# An atypical DYRK kinase connects quorum-sensing with posttranscriptional gene regulation in *Trypanosoma brucei*

**DOI:** 10.1101/771873

**Authors:** Mathieu Cayla, Lindsay McDonald, Paula MacGregor, Keith R. Matthews

## Abstract

The sleeping sickness parasite, *Trypanosoma brucei*, uses quorum sensing (QS) to balance proliferation and transmission potential in the mammal bloodstream. A signal transduction cascade regulates this process, a component of which is a divergent member of the DYRK family of protein kinases, TbDYRK. Phylogenetic and mutational analysis in combination with activity and phenotypic assays revealed that TbDYRK exhibits a pre-activated confirmation and an atypical HxY activation loop motif, unlike DYRK kinases in other eukaryotes. Phosphoproteomic comparison of TbDYRK null mutants with wild type parasites identified molecules that operate on both the inhibitory ‘slender retainer’ and activatory ‘stumpy inducer’ arms of the QS control pathway. One of these molecules, the RNA-regulator TbZC3H20, regulates parasite QS, this being dependent on the integrity of its TbDYRK phosphorylation site. This analysis reveals fundamental differences to conventional DYRK family regulation and links trypanosome environmental sensing, signal transduction and developmental gene expression in a coherent pathway.

## Introduction

Eukaryotic cells respond to their environment via signal transduction cascades whose general structure and features have been intensively studied in model eukaryotes. These enable the adaptation to a changing environment or response to external signals that regulate cellular differentiation and specialisation. Many of these signalling cascades have been dissected in detail, revealing broadly conserved networks that regulate molecular function and interactions in evolutionarily separated eukaryotic groups[1]. Among the most common regulatory mechanisms is through the action of proteins kinases and phosphatases, whose activity controls the reversible phosphorylation of approximately 30% of the proteome of eukaryotes, involving proteins implicated in a wide range of functions, including cell growth and cycle control, cytoskeleton organisation, extracellular signal transmission and cell differentiation. The structure of conventional signalling cascades, however, is not necessarily representative of the diversity of eukaryotic life, with the environmental signalling pathways and regulatory pathways outside commonly studied model eukaryotes being poorly understood.

A tractable model to explore the diversity of eukaryotic signalling networks are kinetoplastid parasites, which separated from the eukaryotic lineage at least 500 million years ago[2]. These organisms encode around 190 protein kinases [3] and are exquisitely sensitive to their environment. This sensitivity to well defined environmental stimuli allows the molecular dissection of signalling pathways that drive the differentiation events that characterise the complex life cycles of these organisms. An exemplar is *Trypanosoma brucei,* a kinetoplastid parasite responsible for Human African Trypanosomiasis (HAT) and Animal African Trypanosomiasis (AAT), that is transmitted through the bite of the tsetse fly. One major environmentally-signalled event for *T. brucei* involves their differentiation in the mammal host from a replicative ‘slender form’ to an arrested and transmission-adapted ‘stumpy form’[4]. This differentiation is triggered by a quorum-sensing (QS)-like mechanism where parasites respond to the accumulation of a stumpy-induction factor [5–7]. Once the signal is received, it is transduced via a non-linear hierarchical signalling pathway [8] comprising at least 30 molecules [9]. This includes signal processing molecules, protein kinases and phosphatases and post-transcriptional gene expression regulators as well as additional proteins of unknown function.

One of the components involved in the differentiation process is a molecule related to the protein kinase Yak sub family [8, 9]. Yak kinases belong to the dual-specificity yak-related kinases (DYRK) family included in the CMGC group, which is over represented in trypanosomatids compared to humans [10]. The DYRK family is subdivided into 5 sub families, the homeodomain-interacting protein kinases (HIPKs), the pre-mRNA processing protein 4 kinases (PRP4s), the Yak kinases (present in lower eukaryotes only), the DYRK1 and the DYRK2 kinases (reviewed by Aranda et al. [11]). In mammals, the DYRK1/2 sub families are characterised by the DYRK-homology (DH)-box upstream of a kinase core that contains the ATP binding domain and the activation loop. The activity of DYRK kinases is dependent on auto-phosphorylation of the second tyrosine residue in the YxY motif present in the activation loop, and they phosphorylate substrates on serine and threonine residues. The full activity of mature DYRK proteins may also require other phosphorylation events outside the kinase core[12] or may depend on their interaction with other proteins [13] [12]. Several mammal DYRKs have also been implicated in cell differentiation. For example, human DYRK1A/ murin DYRK1B and human DYRK2 have been suggested to have a role in differentiation mediated by cell cycle arrest in G1/G0 and G2/M, respectively [14–16].

Here we describe the essential role of several unconventional features in the activity of an atypical DYRK-family kinase identified in the parasite *T. brucei* and that plays a major role in the quorum-sensing stimulated development from slender to stumpy forms in the mammalian bloodstream. We also describe the identification of *T. brucei* DYRK (TbDYRK) substrates and reveal their involvement in both inhibitory and stimulatory arms of the developmental control pathway. This places TbDYRK at a pivotal position in the control of parasite developmental competence, connecting signal transduction to gene expression regulatory processes.

## Results

### Tb927.10.15020 encodes for a divergent kinase of the DYRK family

To initially analyse gene Tb927.10.15020, we performed a phylogenetic analysis using the kinase core of all members of the CGMC kinase family from human, *C. elegans*, *D. melanogaster*, *S. cerevisiae* and all identified members of the DYRK family from *T. brucei* [3]. This identified a DYRK1 family kinase (Tb927.5.1650) and a previously characterised DYRK2 member (TbDYRK2; Tb927.11.3140; [17]). The analysis revealed that Tb927.10.15020 is a divergent DYRK belonging to a paraphyletic group of DYRK2 (Figure 1a). Indeed, the two *C. elegans* kinases with which the trypanosome protein clusters, Ce_C36B7.1 and Ce_C36B7.2, have been identified as DYRK2 kinases in the kinase.com database (http://kinase.com) and as DYRK3 in the wormbase.org database (https://wormbase.org) [18]. It is also notable that Tb927.10.15020 presents a potential divergent DH box (NEx_(2)_DDx_(3)_Y) [19] specific for DYRK1/2, a divergent NAPA-1 region (Lx_(3)_Ex_(2)_Ex_(15)_G), but no NAPA-2, both specific for DYRK2 and present in the previously analysed TbDYRK2 [17, 20] (Figure 1b). Multiple sequence alignment of the kinase core of Tb927.10.15020 with several well characterised members of the DYRK family from a number of model organisms (Figure S1), also revealed the presence of three long inserts in the trypanosome kinase. The kinase is present in the genome of different trypanosomatid organisms and sequences of the first two inserts are relatively well conserved among these orthologues, while the sequence of the last insert in Tb927.10.15020 is poorly conserved outside the *brucei* group (Figure S2). These multiple sequence alignments also revealed the presence of a serine (S856) instead of the classical glycine in the highly conserved DFG motif (Figure 1b, Figure S1), as well as the presence of a histidine (H866), where usually is found a phosphorylable residue, in the HxY motif of the activation loop (Figure 1b, Figure S1). Analysis of this unconventional activation loop in various species revealed the motif was absent in yeast DYRKs, whereas one of 11 human DYRKs present such a motif and one of six in Drosophila. In contrast, of the seven DYRKs identified in the *T. brucei* kinome [3], four present unconventional activation motifs, of which two include a histidine (Tb927.10.15020 and Tb927.5.1650), suggesting a particular mode of regulation in trypanosomes more developed than in other species. Given the association with the DYRK family, we henceforth refer to the protein encoded by Tb927.10.15020 as TbDYRK (previously termed ‘YAK’ in [9] and [8]).

**Figure 1:**
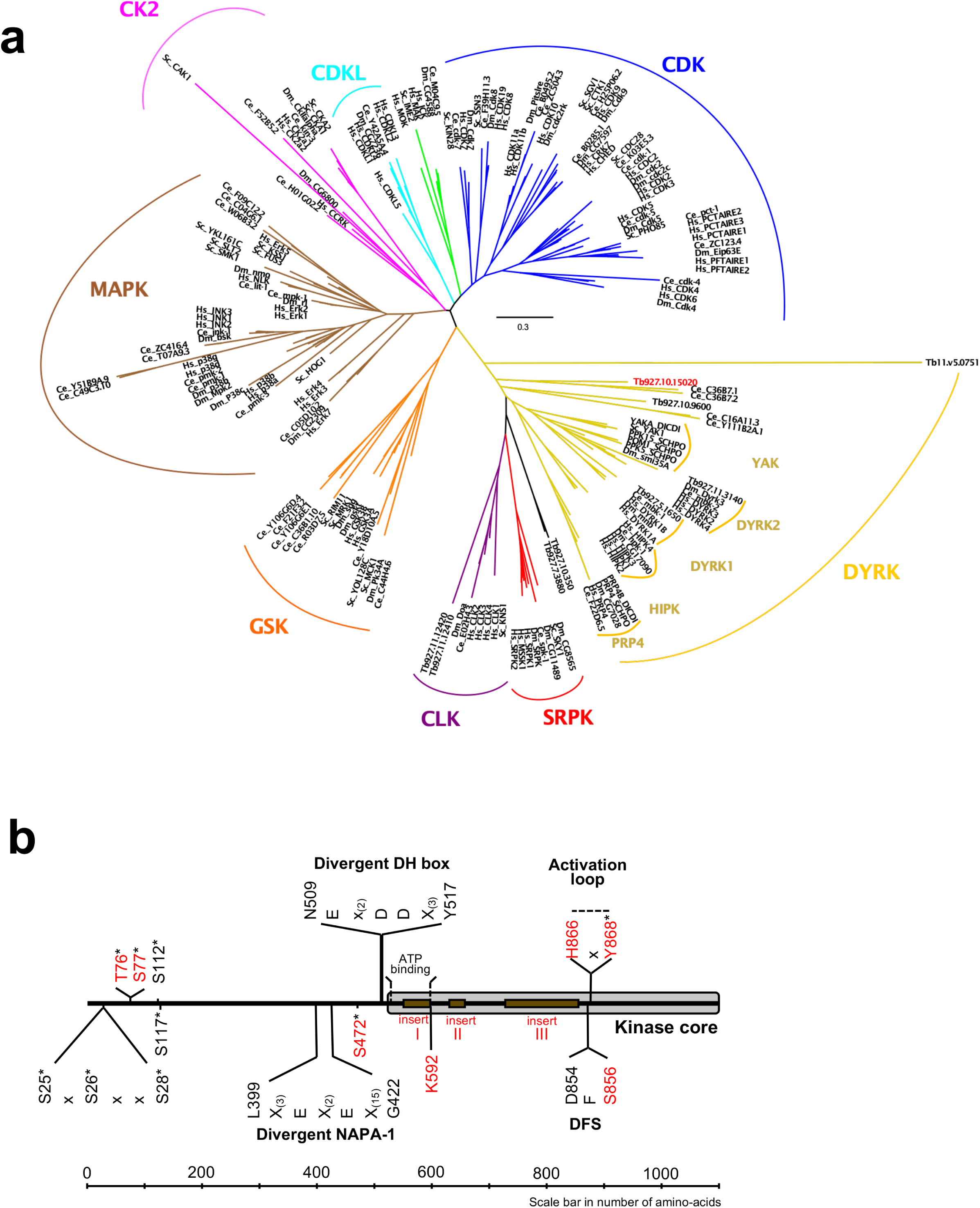
Phylogenetic analysis of the CMGC protein kinase family. **a.** The evolutionary history was inferred by using the Maximum Likelihood method based on the Whelan And Goldman + Freq. model [62]. Initial tree(s) for the heuristic search were obtained by applying the Neighbour-Joining method to a matrix of pairwise distances estimated using a JTT model. The tree is drawn to scale, with branch lengths measured in the number of substitutions per site. All positions with less than 95% site coverage were eliminated. That is, fewer than 5% alignment gaps, missing data, and ambiguous bases were allowed at any position. Hs = *Homo sapiens*, Ce = *Caenorhabditis elegans*, Dm = *Drosophila melanogaster*, Sc = *Saccharomyces cerevisiae*, DICDI = *Dictyostelium discoideum*, Tb927 = *Trypanosoma brucei*. The early divergent TbDYRK is highlighted in red. **b.** Schematic representation of linear protein sequence of TbDYRK, highlighting particular characteristics of its sequence. Identified phosphosites are represented by an *. Insertions I, II and III are presented with the brown boxes. All residues or inserts mutated or deleted in this study are represented in red.

### TbDYRK activity is required for the slender to stumpy differentiation

We initially confirmed the role for TbDYRK in differentiation competence described by [9] and [8]. A wild-type differentiation competent cell line and a previously validated TbDYRK knock-out (KO) cell line [8] were assayed *in vitro* using the cell permeable cyclic AMP analogue, 8-pCPT-cAMP, that promotes the expression of some characteristics of stumpy forms ([21, 22]). As expected, the wild-type cells differentiated in response to treatment, causing a 60% growth inhibition at 96 hours compared to untreated cells, whereas the TbDYRK KO cells had reduced differentiation competence, with only 23% growth inhibition at the same timepoint (Figure 2a). To gain further insight into the specific function of this kinase and the role of selected residues for its kinase activity, we also created two further cell lines ectopically expressing either a TY-YFP-tagged non-mutated version of the kinase (‘NM’) or a predicted ‘kinase dead’ mutant with lysine 592 converted to alanine (‘K592A’); in each case these were expressed in cells retaining the endogenous gene. We then induced ectopic kinase expression using doxycycline and compared the response of each cell line to 8-pCPT-cAMP. Ectopic expression of the NM version of the kinase resulted in an increased differentiation response to 8-pCPT-cAMP compared to wild-type cells (86% growth inhibition at 96 hours), whereas ectopic expression of the K592A mutant considerably reduced differentiation (27% growth inhibition at 96 hours), despite the presence of the endogenous wild-type gene, presumably through a dominant-negative mechanism (Figure 2a).

**Figure 2:**
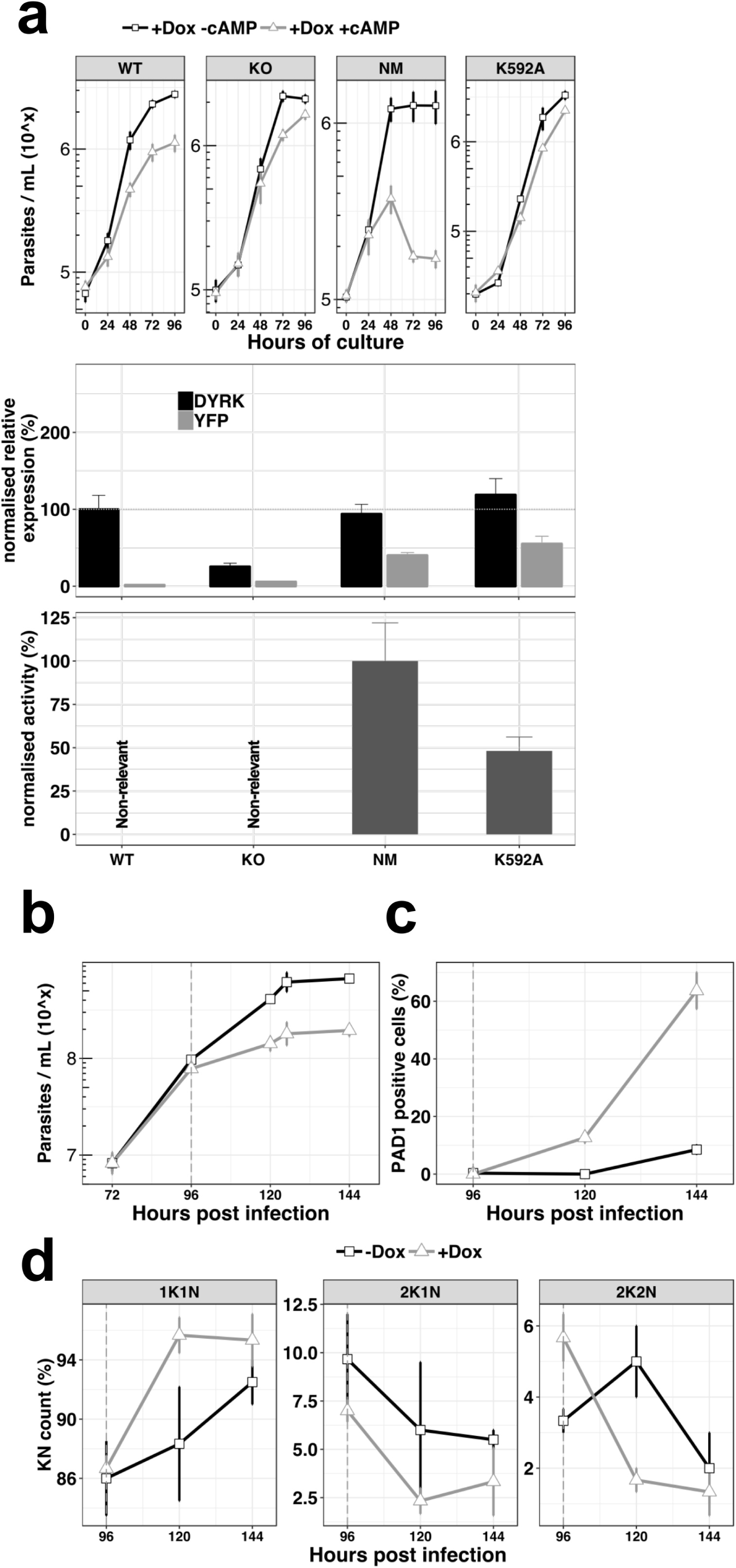
Ectopic expression of the TbDYRK drives stumpy differentiation. **a.** The top panel shows the *in vitro* phenotype analysis after expression of TbDYRK in different strains treated with doxycycline in presence (black) or absence (grey) of pCPT-cAMP (n = at least 2 experiments in 3 replicates, error bars=SD). WT = Parental cell line, KO = Knock out cell line for the gene TbDYRK, NM = Ectopic expression of the non-mutated version of TbDYRK, K592A = Ectopic expression of TbDYRK carrying the mutation K592A. The middle panel represents the mRNA level of expression of TbDYRK (in black), including the endogenous and the ectopic gene when present, and of the YFP tag of the ectopic fusion gene (in grey) (n = 1 experiment in 3 replicates; error bars=SEM). The dotted horizontal grey line represents 100% of expression (mean), obtained from the expression of TbDYRK in the WT cell line after 24h of incubation with doxycycline and no pCPTcAMP. Note that background signal by qRT-PCR is retained with the TbDYRK KO cell line; independent northern blotting has confirmed absence of the transcript. The lower panel represents the activity of the purified kinase against the generic substrate Casp9 as measured by radioactive kinase assay. The mean level of activity of different experiments of NM has been set at 100% of activity (n = at least 2 experiments in 3 replicates; error bars=SEM). **b.** Parasitaemia measured in the bloodstream of mice infected (n=3, error bars=SD) with the NM strain and treated (+Dox, grey) or not (-Dox, black) after 96h post infection. **c.** Percentage of PAD1 positive cells in the same blood smears. **d.** K(inetoplast) N(uclear) scoring from DAPI staining of blood smears slides of the mice infected as previously described. n=250 cells, error bars=SD.

Levels of TbDYRK kinase protein expression are undetectably low, even using epitope tagging. Therefore, as an alternative to monitor the level of expressed kinase, we measured the total amount of TbDYRK mRNA by RT-qPCR, either detecting both the endogenous and the ectopic gene-derived mRNAs or, by specifically targeting the YFP component, only the ectopic gene-derived mRNA (Figure 2a, middle panel). Interestingly, when TbDYRK expression was examined in the NM kinase line, the total amount of TbDYRK transcript did not exceed the level in WT cells (94 ± 12.5 % of expression), despite the ectopically expressed gene contributing 40 ± 3.7% of the total TbDYRK mRNA. These results suggest that the ectopic expression of the NM kinase leads to a reduction of the endogenous mRNA and that the total amount of mRNA may be regulated. With the expression of the K592A mutant, the total amount of mRNA was 119 ± 21.0% compared to WT cells, of which 55 ± 10.0% corresponded to the K592A-YFP fusion.

Next, we determined the activity of the NM and predicted kinase-dead mutant *in vitro*. To identify a suitable substrate for this assay we purified the expressed tagged NM kinase from insect cells and performed an *in vitro* kinase assay in the presence of a panel of candidate generic substrates (Figure S3a). The first 200 amino acids of the *Mus musculus* caspase 9 (Casp9) was the best available substrate, and was used to perform kinetic analysis to determine the optimal conditions to assay TbDYRK activity (Figure S3b and c). We then applied this methodology to measure the phosphotransferase activity of the NM and K592A mutants, also expressed in and purified from insect cells. The mean of the activity measured with the NM was set at 100% and the results indicated that the mutant K592A exhibited reduced activity compared to NM (48.2 ± 8.0 %, Figure 2a, lower panel), demonstrating that the predicted ‘kinase dead’ mutation reduced but did not completely eliminate activity.

To confirm the *in vitro* results suggesting that the ectopic expression of the active NM kinase leads the cells to arrest at a lower concentration and potentially drives the cells to stumpy-like differentiation earlier, we infected mice with the NM strain. The data in Figure 2b demonstrates that the parasitaemia was reduced after induction of the expression of the non-mutated kinase by provision of doxycycline at 96h post infection. Further, the cells presented a strong increase of PAD1 expression as assessed by IFA (Figure 2c) (63.7 ± 11.0 % versus 8.5 ± 2.1 % at 144h post infection, uninduced and induced respectively) and arrested in G1/G0 earlier (Figure 2d). These results show that the ectopic expression of active TbDYRK drives the cells to differentiate to stumpy forms at a lower parasitaemia.

### N-terminal phosphosite-residues and the HxY motif of the activation loop are essential for TbDYRK activity

We next analysed the role of selected residues and domains of the TbDYRK molecule to explore their difference from conventional eukaryotic DYRKs. First, we and others have identified nine phospho-sites on the TbDYRK protein (asterisks on Figure 1b)[8, 23, 24]. Eight of these are residues located in the N-terminal domain of the protein, and one, Y868, belongs to the atypical HxY motif of the activation loop. We assessed the role of three of the N-terminal residues for kinase function, i.e., T76, S77 and S472, by mutating them to alanine and ectopically expressing the mutants in cells retaining the endogenous gene copies. Thereafter, similar analyses to those previously were used, namely i) scoring cell growth upon ectopic expression of the mutants in the presence or absence of 8-pCPT-cAMP in vitro (Figure 3a), ii) quantification of the mRNA level of each mutant and total TbDYRK (Figure 3b), and iii) quantification of the phospho-transferase activity of each protein expressed in insect cells (Figure 3c). The results obtained for the WT, KO and NM repeat those presented in Figure 2; the remainder are a combination of at least 3 independent experiments in 3 replicates for each mutant. Expression of the T76A and S77A mutants did not change the response to 8-pCPT-cAMP of the cells compared to WT parasites (Figure 3a). Both mutations render the kinase completely inactive (Figure 3c) and their mRNAs were well expressed (Figure 3b). For both mutations, mRNAs derived from the mutated ectopic copy represented the majority of the total TbDYRK mRNA in each cell line (68.9 ±13.35, 71.46 ± 16.52%) although the combined mRNA level was not elevated over wild type abundance, implying that expression of the mutants causes regulation of transcripts derived from the endogenous gene as seen earlier. The mutation of S472, by an alanine, also completely disrupted the kinase activity and, similar to the phenotype observed with the KO cell line, generated resistance to 8-pCPT-cAMP treatment, suggesting the inactive expressed kinase prevented the function of TbDYRK expressed from the endogenous locus (Figure 3). This phenotype could be contributed to by the high level of expression of the tagged gene, with the combined endogenous and ectopically expressed TbDYRK mRNA reaching 173.2 ± 29.82% of TbDYRK in wild type cells.

**Figure 3:**
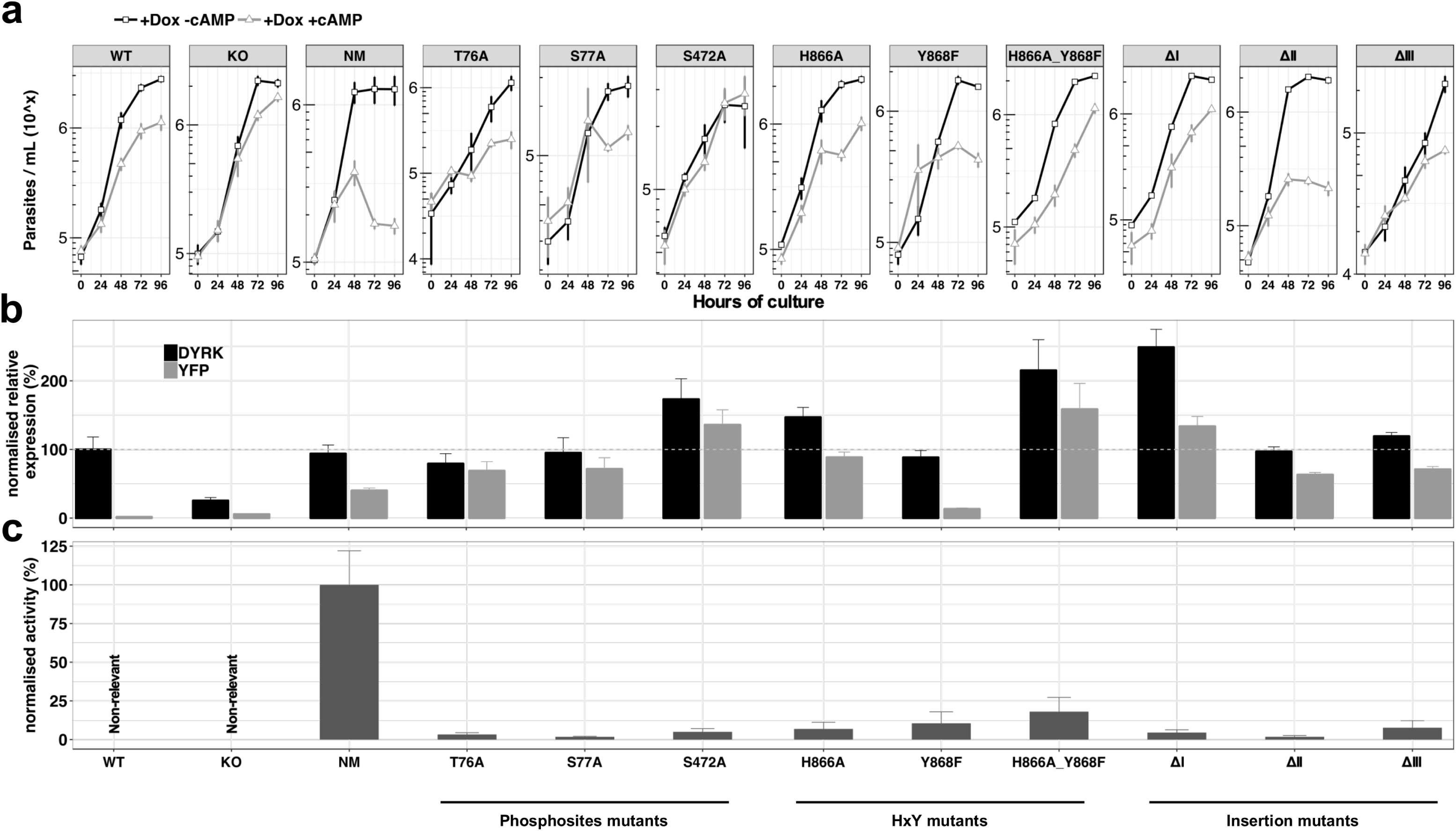
Analysis of the unconventional features of TbDYRK with respect to their phenotype, expression and activity. **a.** *In vitro* phenotype analysis after expression of TbDYRK mutants treated with doxycycline in the presence (black) or absence (grey) of pCPT-cAMP (n = at least 2 experiments in 3 replicates; error bars=SD). WT = Parental cell line, KO = Knock out cell line for the gene TbDYRK, NM = Ectopic expression of the non-mutated version of TbDYRK. WT, KO and NM results are the same as presented in Figure 2 and are used here as control; other columns reflect the respective mutants analysed **b.** mRNA level of expression of TbDYRK (in black), including the endogenous and the ectopic gene when present, and of the YFP tag of the ectopic fusion gene (in grey) (n = 1 experiment in 3 replicates, error bars=SEM). The dotted horizontal grey line represents 100% of expression (mean), obtained from the expression of TbDYRK in the WT cell line after 24h of incubation with doxycycline and no pCPT-cAMP. **c.** Activity of the purified kinase against the generic substrate Casp9 as measured by radioactive kinase assay. The mean level of activity of different experiments of NM has been set at 100% of activity (n = at least 2 experiments in 3 replicates; error bars=SEM).

We next investigated the function of the unconventional HxY motif in the activation loop, where a phosphorylable residue (Y or T) is usually observed instead of the histidine residue in TbDYRK. As previously, we analysed the effect of the mutations H866A, Y868F and the combination H866A_Y868F on the response to 8-pCPT-cAMP treatment, on TbDYRK mRNA levels and on kinase activity (Figure 3a-c). Mutations of these residues disrupt the kinase activity with an activity level of 6.84 ± 4.34 % (H866A), 10.54 ± 7.4 % (Y868F) and 18.02 ± 9.25 % (H866A_Y868F) compared to non-mutated (NM) protein. Expression of mutant H866A led to a similar response as in WT cells to 8-pCPT-cAMP. The TbDYRK Y868F mutant mRNA was expressed at a lower level than the NM mRNA (13.24 ± 1.3%), and the cells responded to 8-pCPT-cAMP similarly to WT cells. In contrast, expression of a mutant with both mutations (H866A_Y868F) led to a slight resistance to 8-pCPT-cAMP, with the mutant being highly expressed and comprising the majority of TbDYRK mRNA in the cell, with reduction in the contribution of the mRNA from the endogenous TbDYRK allele, suggesting a negative regulation.

Overall, these data show that mutation of any of the four putative phosphosite-residues, either in the N-terminal domain or the HxY motif in the activation loop, disrupt TbDYRK kinase activity. Despite the different mutations having the same apparent effect of creating kinase-dead mutants, a range of phenotypes (increased, decreased or unchanging responsiveness to 8-pCPT-cAMP) were observed when these were ectopically expressed in cells.

### The three inserts inside the kinase core have a major role in the kinase function

As highlighted earlier, the conventional kinase core sequence of TbDYRK is interrupted by three inserts of 36, 22 and 119 amino-acids (Figure S1), which are predicted to be positioned peripherally to the core enzyme structure and so amenable to functional analysis by deletion (Figure S4a and b). Each insert was, therefore, deleted and the expressed mutants assayed for their response to 8-pCPT-cAMP (Figure 3; ΔI, II or III). Activity assays demonstrated that each of the three inserts was essential for the phospho-transferase activity of the kinase. The ΔI mutant was highly expressed, and phenotypically, this expression rendered the cells relatively resistant to 8-pCPT-cAMP-induced arrest despite the presence of the endogenous TbDYRK allele. The ΔII mutant was expressed at a similar level as NM; however, despite this and the lack of activity of this mutant, a similar phenotype as the expression of NM was observed. Thus, there was a decrease of the parasite density in response to 8-pCPT-cAMP treatment that was greater than seen with WT cells. In contrast, the inactive ΔIII mutant generated a similar phenotype to WT cells in response to the 8-pCPT-cAMP, despite its effective expression.

To explore the phenotype of the expression of the ΔII mutant, which matched the ectopic expression of the NM kinase and yet lacked kinase activity we performed a viability assay of cells where the NM or the ΔII mutant were expressed in the presence or absence of 8-pCPT-cAMP (Figure S5). This revealed that expression of the inactive ΔII mutant resulted in cell death after 8-pCPT-cAMP exposure whereas those expressing the NM kinase were viable. Apparently therefore, expression of the ΔII mutant is toxic to cells exposed to 8-pCPT-cAMP but not those in the absence of this stimulus.

### The essential S856 in the DFS motif allows a potential pre-activation state

We next investigated the role of S856 that contributes to the unconventional DFS motif of TbDYRK. This motif, usually comprising DFG, functions in the conformational changes upon kinase activation necessary for the correct alignment of the catalytic residues and the binding of ATP. The mobility brought about by the glycine residue allows the switch of the phenylalanine from the inactive state ‘DFG-out’ to the active state ‘DFG-in’ [25].

By analysis of the *T. brucei* kinome, we firstly established that unconventional DFG motifs (either absent or mutated on the phenylalanine and/or the glycine) were present on 18.5% of *T.brucei* ePKs, and that 8 kinases possess a change of the glycine to either an aspartic acid, an asparagine or a serine (Figure 4a). We note that the DFS motif is present on four kinases in *T. brucei* (Tb927.9.1670, Tb927.1015020, Tb927.10.16160 and Tb927.11.5340), representing 2.5% of *T. brucei* ePKs, while it is present in only 0.4% of the ePKs in humans.

**Figure 4:**
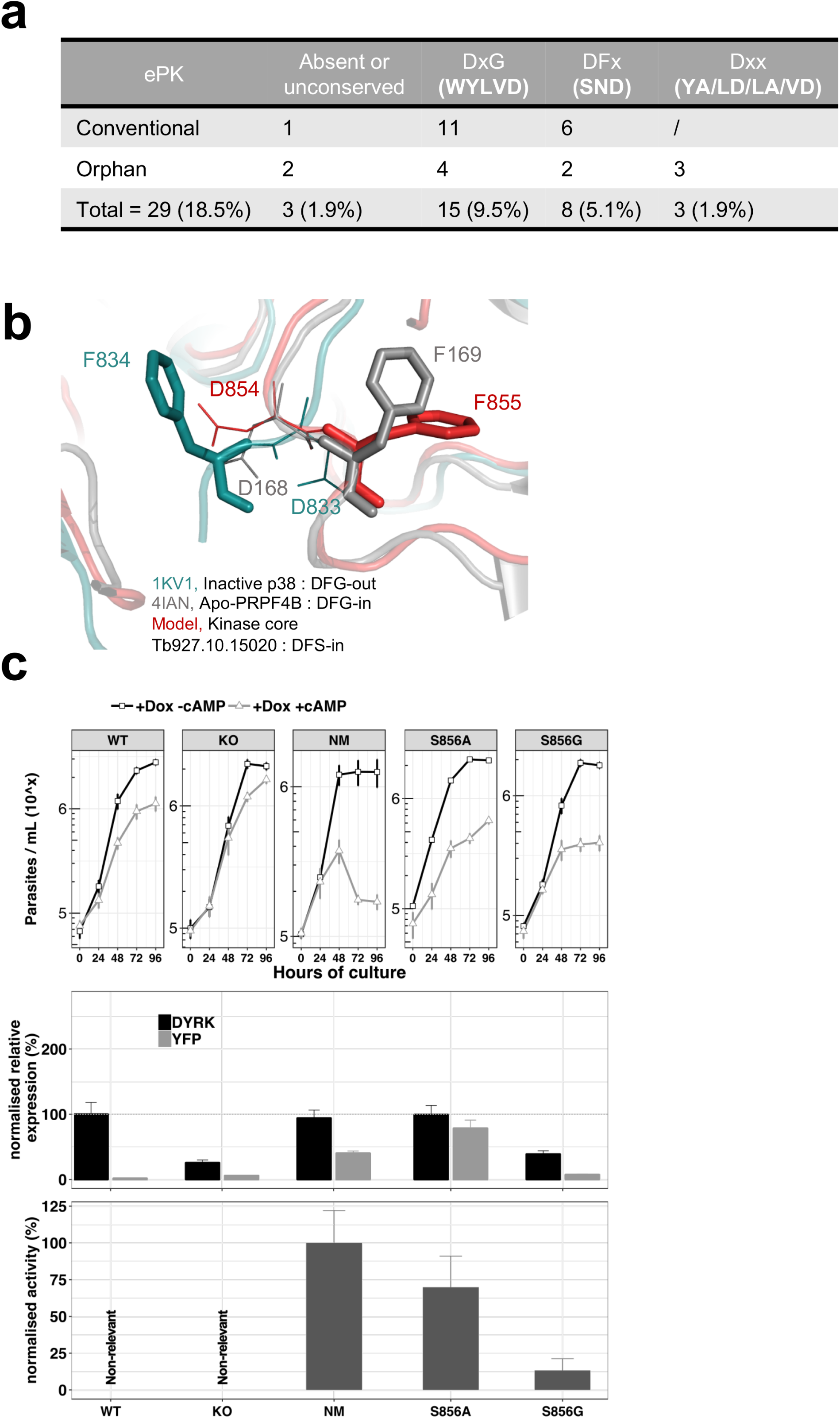
The unconventional DSF motif renders the kinase more rigid and is essential for the activity of the kinase. **a.** Table indicating the unconventional DFG motif in *T. brucei* ePKs. Conventional = conventional kinases, Orphan = orphan kinases. **b.** Model of the TbDYRK kinase core generated by the i-TASSER server, indicating the position ‘DFS-in’ of the DFS motif of the kinase. **c.** Structure/function analysis, as presented earlier, of the DFS motif. WT, KO and NM results are the same as presented in Figure 2 and are used here as control. S856A = Ectopic expression of TbDYRK carrying the mutation S856A, S856G = Ectopic expression of TbDYRK carrying the mutation S856G. For the growth curves, n=3, error bars=SD; for the mRNA and kinase activity assays, n=3, error bars=SEM.

To investigate the role of this serine in the activation state of the kinase, we modelled the kinase core of TbDYRK using the i-TASSER server. The model obtained revealed interesting features, although the overall scores highlighted complexity of the predictions (c-score = −2.73, estimated TM-score = 0.40±0.13, estimated RMSD = 14.5±3.7 Å), undoubtedly influenced by the three inserts generating a more important b-factor due to the consequent low alignment coverage and the absence of predicted secondary structure (Figure S4a). Based on the structural prediction, the N-terminal lobe appears to be well conserved with the presence of the classical beta sheet β1-5 and the αC helix. The C-terminal lobe is composed of the classical beta sheet β6/7 after the hinge region, the α-helix C/D/E/F/G/H and I, and also contains the MK1 insert specific of the CMGC family and the α-helix 7 and 8 [26–28] (Figure S4c). From this analysis we predicted that the presence of this serine in place of glycine leads to a pre-activation state of the kinase with a phenylalanine in position ‘DFS-in’, and the aspartic acid pointing toward the active site, as observed on the crystal structure of active p38 (Figure 4b). In addition, and in support of the hypothesis of a pre-active conformation of TbDYRK, the model suggests a close conformation of the ATP binding pocket with the glutamine 607 at 3.5 Å from the lysine 592, in concordance with the 3.8 Å measured between the corresponding amino-acid of the active structure of p38 (Figure S4c). This observation is also accompanied by fully aligned regulatory and catalytic (R- and C-) spines as observed in active kinases [25, 29, 30] (Figure S4d).

Interestingly, mutation of the serine S856 to alanine (DFA) did not greatly reduce kinase activity (69.82 ± 21.17 %, Figure 4c, lower panel), whereas the mutation to glycine, to generate a classical DFG motif, almost completely disrupted the kinase activity (13.26 ± 7.96), suggesting that the greater mobility brought by the glycine is detrimental for activity. In addition, despite the strong expression of the S856A mutant (Figure 4c, middle panel) and the maintenance of around 70% of its kinase activity in vitro, the same phenotype as WT cell was observed in response to 8-pCPT-cAMP (Figure 4c, top panel). These results suggest either that the remaining activity of this mutant is not sufficient to generate the strong NM phenotype, or that the activity against endogenous substrates may be different to activity against *Casp9* in vitro.

In combination these structural predictions and functional studies demonstrated that phosphorylation at the N terminus of TbDYRK, the presence of atypical inserts and integrity of the activation loop in the kinase core region, and the presence of the unconventional DFS motif all are important for the activity of the atypical kinase leading to a range of phenotypic outputs in response to 8-pCPT-cAMP. Moreover, there is evidence of mRNA regulation when the mutants were responsive to 8-pCPT-cAMP but less when this response is reduced.

### Identification of substrates of TbDYRK implicated in the stumpy differentiation

The next step of the analysis was to explore the signalling pathway in which TbDYRK functions. Therefore, in a first screen, we examined phosphoproteomic changes upon deletion of the TbDYRK gene in slender form parasites. Parental *T*. *brucei* EATRO 1125 AnTat 1.1 90:13 and TbDYRK KO cells were cultured in duplicate at equivalent cell density *in vitro* and protein extracted and analysed after isobaric tandem mass tagging (Flow chart in Figure 5a) as described in [8]. A total of 2499 unique proteins and 7293 unique phosphopeptides were identified; correlations between the replicates were >99% at the peptide level, and at the phosphopeptide level were 0.8805 (*T*. *brucei* EATRO 1125 AnTat1.1 90:13) and 0.9896 (TbDYRK null mutant) respectively, demonstrating excellent reproducibility (Figure S6). The datasets were filtered for peptides with >1.5-fold change in phosphorylation regardless of direction, with an adjusted P value of <0.05, (Supplementary Table 2), revealing that 213 peptides on 172 different proteins were less phosphorylated, and that 191 peptides on 149 unique proteins were more phosphorylated in the TbDYRK KO cell line. As expected, the most depleted protein was TbDYRK itself (Tb927.10.15020). Also, supporting the involvement of the TbDYRK protein in the stumpy differentiation pathway, two sites were identified as less phosphorylated in the null mutant line on Protein Associated with Differentiation (PAD) PAD2 (Tb927.7.5940 - S324: Log2–1.276 - adj P = 0.0128; S309: Log2–1.268 - adj P = 0.0055, Figure 5a, supplementary table S2). In addition, the dataset revealed a strong enrichment of proteins implicated in the regulation of gene expression, posttranscriptional regulation, RNA regulation, phosphorylation and protein and vesicle transport. These data indicate that the absence of TbDYRK influences a broad spectrum of substrates implicated in several essential biological processes, particularly those having a direct action on effectors of gene expression such as the eukaryotic translation initiation factor 4e (Tb927.11.11770), the transcription elongation factor s-II, putative (TFIIS2-1, Tb927.2.3580) or the negative regulator of transcription NOT5 protein (Tb927.3.1920, Figure 5a), for example.

**Figure 5:**
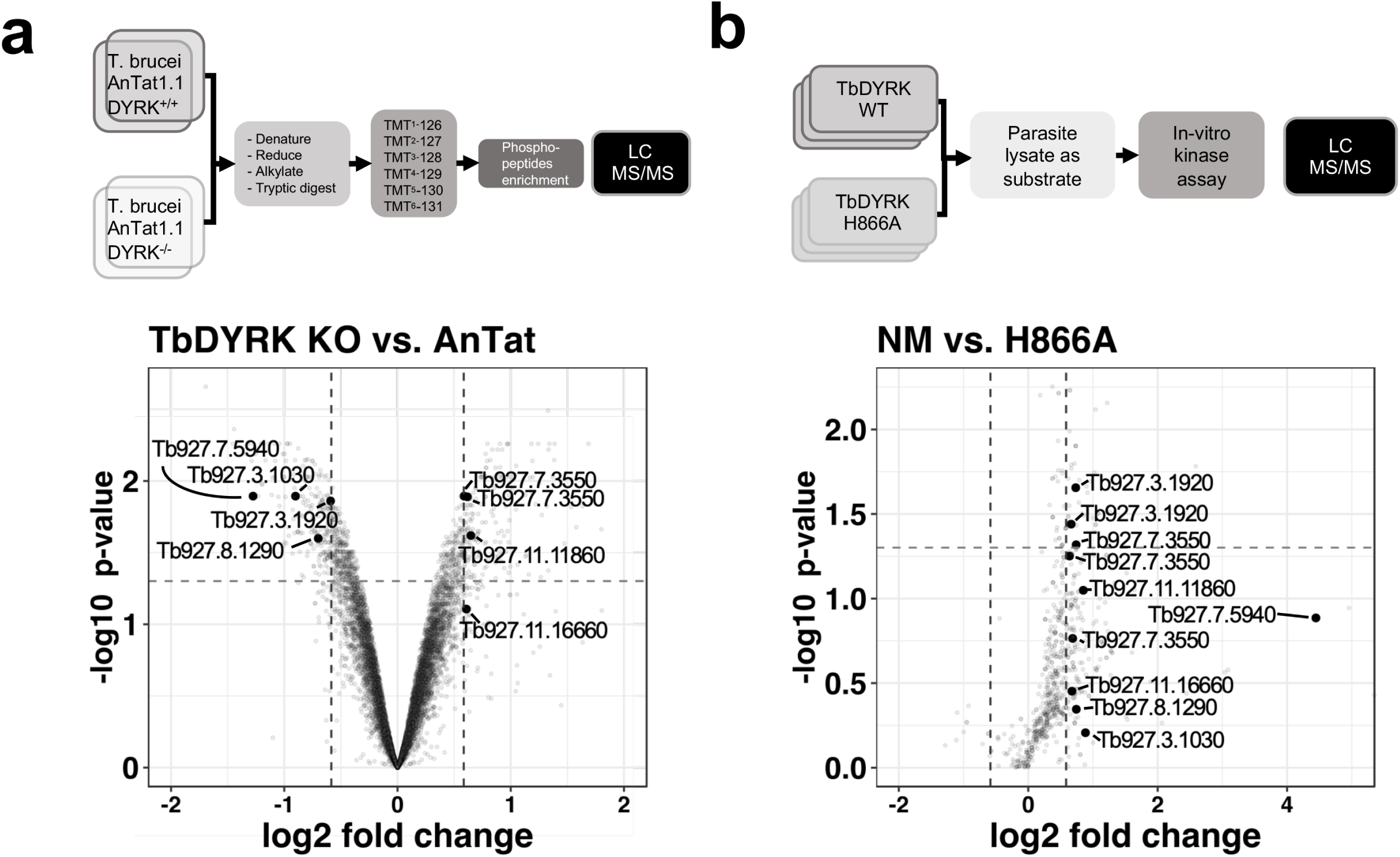
Phosphoproteomic analysis for the identification of substrates of the TbDYRK. **a.** Quantitative phosphoproteomic analysis by TMT isobaric tagging comparing the proteome of WT cells (*T. brucei* AnTat1.1 DYRK+/+) or cells deleted for TbDYRK (DYRK-/-). Top panel: flow chart. Lower panel: Volcano plot of the phosphopeptides with the log2 of the fold change (FC) on the x-axis and the -log10 of the p-value on the y-axis. Vertical dotted lines indicate a |FC|>1.5 and the horizontal dotted line a p-value<0.05. **b.** Quantitative phosphoproteomic analysis comparing the phosphoproteins of cell lysates incubated with the purified active kinase (NM) or inactive (H866A). Top panel: flow chart. Lower panel: Volcano plot of the phosphopeptides with the log2 of the fold change (FC) on the x-axis and the -log10 of the p-value on the y-axis. Vertical dotted lines indicate a |FC|>1.5 and the horizontal dotted line a p-value<0.05. Common peptides in both datasets, presenting a differential phosphorylation >1.5 times disregarding the sense of regulation, are highlighted in both volcano plots.

To filter the extensive list of potential substrates, we took advantage of the structure/function analysis performed previously and used the purified kinase, either as the active (NM) or inactive (H866A) form, to differentially phosphorylate triplicate *T. brucei* procyclic cell lysates previously treated with phosphatase to remove endogenous phosphorylation (Figure 5b). Lysates of this life cycle stage were used to provide sufficient material for analysis. In this dataset, 38 phophopeptides on 29 unique proteins were significantly phosphorylated (FC>1.5 – P-value < 0.05) with the active kinase NM whereas, no significant phosphorylation events were identified with the inactive mutant H866A. As for the previous phosphoproteomic analysis, a GO enrichment of similar functions was also observed in the identified phosphosubstrates i.e. regulation of gene transcription, transport and cellular reorganisation (Supplementary table 3). Comparison of both phosphoproteomic analyses (i.e. whole phosphoproteome and the lysate phosphorylation analysis) demonstrated the enrichment of a kinase substrate consensus motif (R-P/R-x-S/T-P) similar to that of mammalian DYRK 2 and 3 (Figure S7). Overall, 45 proteins were present in both datasets with a |FC|>1.5 (Supplementary table 4) and eight phosphosites were common for seven proteins (|FC|>1.5) with two sites being significant with a P value<0.05 (Figure 5). These two sites were i) serine 395 of the NOT5 protein, which is less phosphorylated in the KO cell line (Log2FC -0.5910 – P value 0.0138) and phosphorylated by the NM kinase (Log2FC 0.7336 – P value 0.0221), and ii) serine 1048 of the WCB cytoskeleton associated protein Tb927.7.3550, which was phosphorylated in the KO cell line (Log2FC 0.5879 – P value 0.0128) and phosphorylated by the NM kinase (Log2FC 0.7402 – P value 0.0481) (Figure 5a, b, supplementary table 4).

We then selected five genes from the 45 proteins present in both datasets with a |FC|>1.5 to individually assess their involvement in the TbDYRK-regulated stumpy formation. These were two hypothetical proteins (Tb927.1.4280 implicated in post-transcriptional activation [31] and Tb927.4.2750), the NOT5 protein previously implicated in the negative regulation of mRNA stability (Tb927.3.1920) [31–35], a zinc finger protein ZC3H2O (Tb927.7.2660) previously shown to be associated to RBP10 protein, to interact with MKT1 and to be implicated in increase of protein translation or in mRNA stability [31, 33], and a bilobe region protein (Tb927.11.15140). Of the genes analysed, we were only able to generate a null mutant cell line for the ZC3H20 protein; for all others, conditional RNAi knock-down cell lines were produced. We then investigated the capacity of the cells to differentiate in response to cell permeable 8-pCPT-cAMP or to stumpy induction factor *in vivo*. Phenotypic analysis of the ZC3H20 KO cell line *in vitro* and *in vivo* demonstrated that this cell line was unable to differentiate into stumpy forms, generating high parasitaemia in mice (Figure 6a), with no PAD1 positive cells detected 144h post infection (Figure 6b) and an absence of cell arrest (Figure 6c), contrasting with the development of the parental line. For the other selected targets, we generated conditional knock-down of the corresponding coding genes by RNAi, and for 2 of them, the hypothetical Tb927.1.4280 (Figure S8) and the bilobe region protein Tb927.11.15140 (Figure S9), did not observe a significant effect on the slender to stumpy differentiation *in vivo*. However, knock-down of hypothetical protein Tb927.4.2750 increased the capacity of the cells to differentiate into stumpy-like cells as judged by their increased sensitivity to the 8-pCPT-cAMP (Figure S10), whereas NOT5 depletion accelerated the capacity to generate PAD1-positive cells in mice (Figure S11). These results suggest that these 2 proteins are “slender retainers” that need to be inactivated or degraded to allow the cells to efficiently differentiate into stumpy forms. In contrast, ZC3H20 is a stumpy inducer, whose activity is required for differentiation. In combination, this reveals a role for TbDYRK on both the inhibitory and stimulatory arms of the stumpy formation pathway.

**Figure 6:**
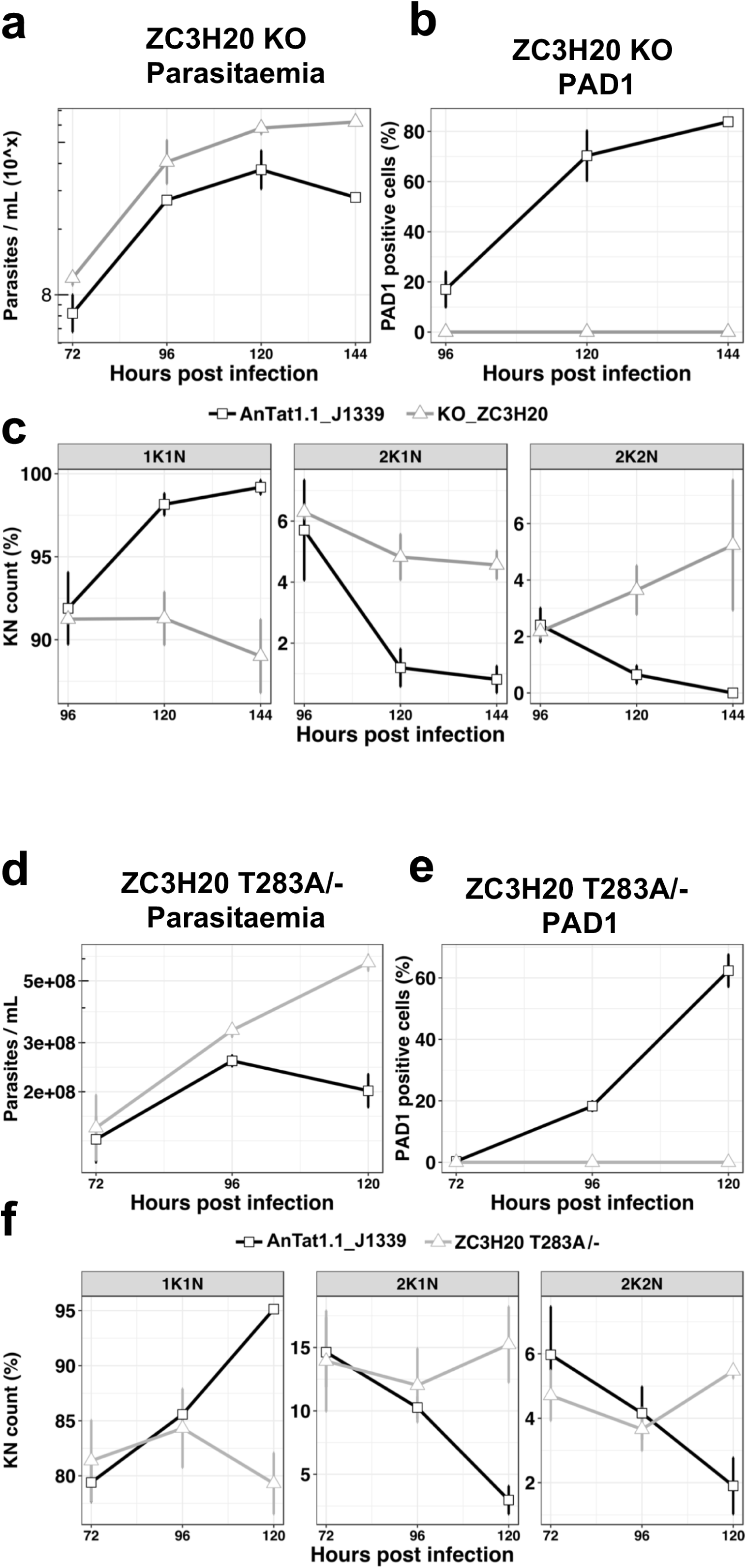
The TbDYRK substrate TbZC3H20 is implicated in the slender to stumpy differentiation. a. Parasitaemia of the ZC3H20 gene KO line (KO_ZC3H20) compared to the parental cell line (AnTat1.1_J1339), n=3, error bars=SD b. Percentage of PAD1 positives cells of the ZC3H20 gene KO line (KO_ ZC3H20) compared to the parental cell line (AnTat1.1_ J1339) on different days of in vivo infection, n=3, error bars=SD c. Percentage of 1K1N, 2K1N and 2K2N cells on different days of *in vivo* infection. n=3, error bars=SD. d. Parasitaemia of the ZC3H20 T283A/-line compared to the parental cell line (AnTat1.1_ J1339), n=3, error bars=SD e. Percentage of PAD1 positives cells of the ZC3H20 T283A/-line compared to the parental cell line (AnTat1.1_ J1339), n=3, error bars=SD f. Percentage of 1K1N, 2K1N and 2K2N cells of the ZC3H20 T283A/-line compared to the parental cell line (AnTat1.1_ J1339) on different days of in vivo infection, n=3, error bars=SD.

### TbDYRK phosphosite mutation on TbZC3H20 abolishes differentiation competence

To confirm the role of TbDYRK in the regulation of the developmental pathway we used CRISPR mediated allelic mutation to replace the identified TbDYRK phosphosite on TbZC3H20 with a non phosphorylable residue in *T. brucei* EATRO 1125 AnTat1.1 J1339 cells[7]. A cell line was successfully generated in which one allele of the TbZC3H20 was replaced by the phosphosite mutant (TbZCH20^T283A^) and the other one deleted (Figure S12a and b); this cell line was predicted to be unresponsive to TbDYRK mediated phosphorylation and hence unresponsive to the QS signal. Growth of the TbZCH20^T283A/-^ cell line *in vivo* in comparison with parental phospho-competent TbZCH20^+/+^ cell lines demonstrated that the mutants were hypervirulent in mice, with the parasites retaining a slender morphology, contrasting with the wild type parasites or a cell line containing one mutated allele and one WT allele (TbZCH20^T283A/+^; Figure S12c), which became stumpy from day 5 of infection (respectively Figure 6d and S12d). The TbZCH20^T283A/-^ mutant parasites also did not express PAD1 (Figure 6e) and did not exhibit cell cycle arrest in G1/G0 (Figure 6f), confirming their developmental incompetence. This was not related to overall mRNA levels; the ZCH20 mRNA levels in the TbZCH20^T283A/-^ line were approximately equivalent to a distinct single allele replacement line that is differentiation competent (Figure S12b). Thus, ZC3H20 is a substrate of the TbDYRK kinase that positively regulates differentiation through its phosphorylation potential, directly linking signal transduction mediated through TbDYRK to an effector of the differentiation pathway with post-transcriptional regulatory activity.

## Discussion

The quorum sensing signalling pathway of trypanosomes in their mammalian host provides a framework for evolutionary analysis of signalling in a group separated from the major eukaryotic models[2]. Here we have analysed through sequence-guided functional analysis and phospho-substrate identification an atypical DYRK component representative of a widely conserved family of proteins centrally implicated in developmental control. This has revealed striking divergence from precedent for this kinase group, particularly an atypical activation mechanism involving unconventional DFG and HxY motifs. It has also defined the first regulatory cascade directly linking environmental sensing, signal transduction and post transcriptional regulation in a kinetoplastid parasite.

In other organisms, from yeast to mammals, DYRK kinases have roles, among other cellular functions, in cell differentiation [11], stress responses, neural development, myoblast development and embryogenesis, with perturbed DYRK family activity implicated in downs syndrome and some cancers. The involvement of TbDYRK in the control of the trypanosome development, therefore, highlights the evolutionarily conserved involvement of this kinase family in differentiation processes across the diversity of eukaryotic life. Despite this, TbDYRK possesses unique characteristics that suggest kinetoplastid specific features of this kinase distinct from other eukaryotes. Particularly, our analysis of TbDYRK has generated the following model for the activation/function of this molecule (Figure 7a). In this model, the kinase would be expressed in slender forms and auto-phosphorylated, possibly in *trans*, as previously suggested through the action of the divergent NAPA-1 domain [17], on the Y868 of the activation loop during translation, as is typical for this kinase family [20, 36]. This auto-phosphorylation at the activation loop, the lack of a second phosphorylable residue on the activation loop (with H in place of tyrosine) and the essential role of the serine the DFS motif (replacing the conventional DFG motif) which is predicted to increase rigidity, suggest this kinase is preserved in a pre-activation state. Indeed, the mutation of this serine to alanine (DFS>DFA) maintained ∼70% of kinase activity *in vitro,* unlike mutation to a glycine which significantly reduced kinase activity. In addition, the modelling of the kinase core shows that the DFS motif is in position ‘DFS-in’, supporting the probable pre-activation state of the kinase, that would require other mechanisms for activation. For example, the kinase could be activated by phosphorylation on at least 3 sites in the N-terminal domain (T76, S77, S472) in response to stumpy induction factor *in vivo* or in response to cell permeable 8-pCPT-cAMP *in vitro*. The functional analysis also revealed the presence of 3 inserts in the kinase core of the protein that are essential for kinase function. These may assist binding to partner proteins [37, 38], similar to human DYRK2, that also acts as an adaptor for the formation of the E3 ubiquitin ligase EDPV [14]. Indeed, the deletion of Insert II, which is cytocidal when cells are exposed to 8-pCPT-cAMP treatment might be caused by effects on the capacity for binding partner interaction.

**Figure 7:**
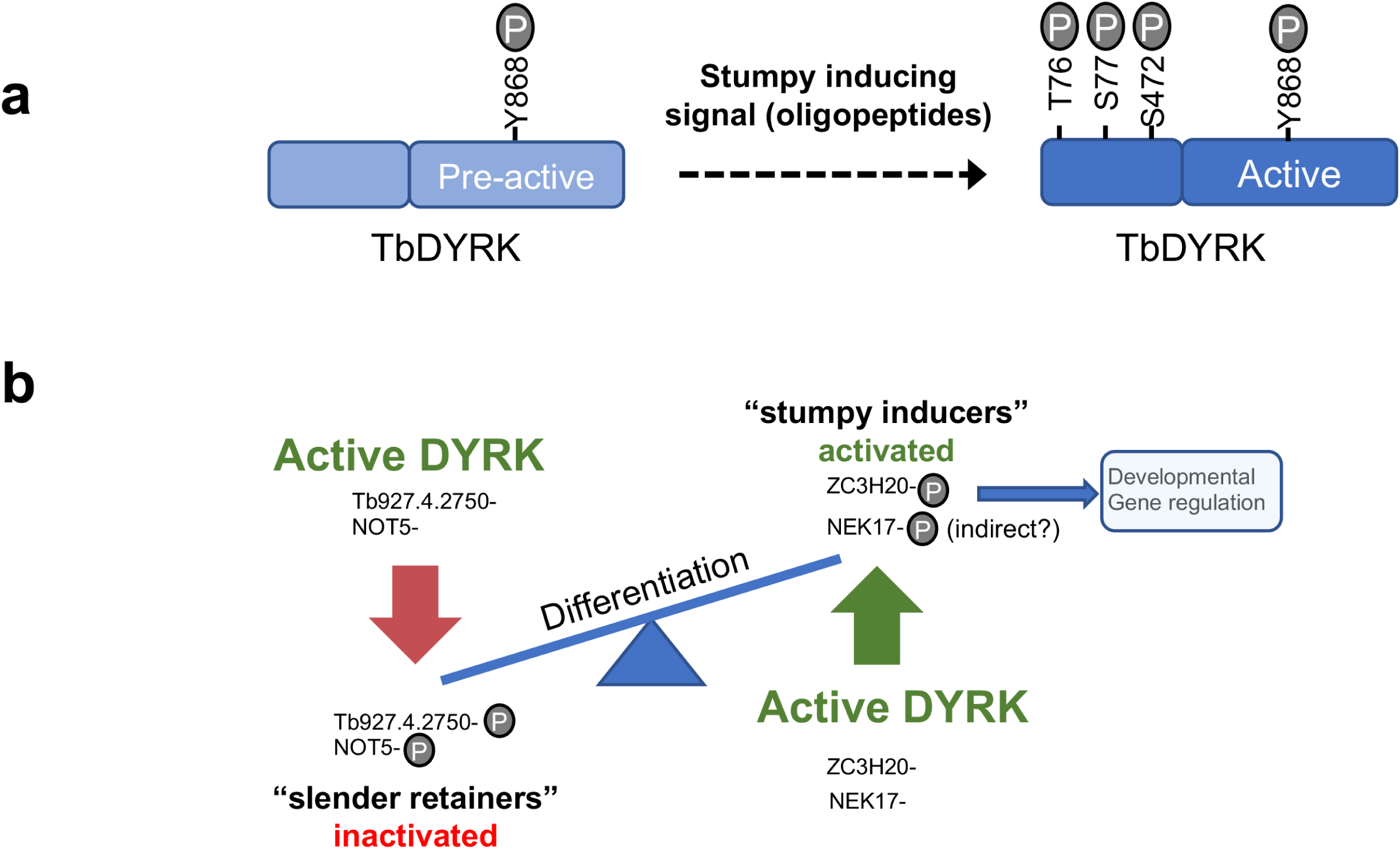
Model of the activation mechanisms and function of the TbDYRK. a. Phosphorylation of the pre-active kinase in response to the stumpy inducing signal results in activation of the kinase. b. Consequences of the activation of TbDYRK for the regulation of differentiation, through the inactivation of slender retainer molecules and the activation of stumpy inducers.

We also observed that the ectopic expression of TbDYRK influenced the total level of TbDYRK mRNA derived from the ectopic and the endogenous derived mRNAs, with this control being relaxed when cells exhibited reduced differentiation. The regulatory effect was not fully correlated with the kinase activity of the expressed TbDYRK mutants but did correspond to the differentiation phenotype generated by the expression of the mutants and so may relate to the activity of other components in the pathway. The combination of kinase activity and/or protein-protein interactions of TbDYRK may control the activity of post transcriptional regulators, which would then promote differentiation through regulated mRNA stability or translation of downstream targets, but also regulate the level of TbDYRK mRNA itself. Stringent regulation of TbDYRK levels would prevent the parasite irreversibly differentiating to stumpy forms prematurely and is reflected in the very low levels of TbDYRK mRNA and protein detectable in bloodstream forms.

Two complementary quantitative phosphoproteomic approaches to identify substrates of the kinase were then performed. First, we compared the phosphoproteome of WT and KO cells and identified around 400 peptides significantly differentially regulated between the cell lines. Interestingly, we observed an enrichment of proteins implicated in chromatin reorganisation and transcriptional regulation, mRNA stability and translation. For example, the histone deacetylase HDAC3 (Tb927.2.2190) presented reduced phosphorylation in the KO cell line, and the mammal DYRK1B has been shown to phosphorylate HDAC5 and 9 to promote myocyte differentiation [39]. A previous study, on the hierarchical organisation of components implicated in the signal transduction of the stumpy differentiation pathway, has identified potential substrates of the MEKK1 kinase, another protein implicated in this process [8]. Comparing the datasets, 12 proteins were shared between the null mutants for each kinase that were less phosphorylated, including the RNA-binding protein, RBP31 (Tb927.4.4230), which has been implicated in decreased translation or mRNA stability [32, 40], kinetoplastid kinetochore KKT4 protein (Tb927.8.3680) that is essential for chromosomal segregation [41] and NEK17 (Tb927.10.5950), another component of the QS signalling pathway [9]. The latter protein exhibits reduced phosphorylation on threonine 195 in both datasets, suggesting that NEK17 phosphorylation requires both TbDYRK and MEKK1 either directly or through the action of another kinase.

Phosphorylation of parasite lysate substrates with the purified active kinase identified 29 proteins that were significantly phosphorylated by the active kinase compared to the inactive mutant. Observed differences between the phosphoproteomic analysis of the null mutant and the lysate analysis are likely due to the technical and biological reasons. Firstly, the use of isobaric tags for analysis of the null and wild type parasites is more sensitive. Secondly, direct and indirect phosphorylation events could contribute to the differences observed. DYRKs are known to act as priming kinases, such that their phosphorylation of substrates is required for further phosphorylation by other kinases, such as GSK3 or PLK [42–44]. This could contribute to the reduction of phosphorylated proteins detected in the null mutant line that are not directly phosphorylated by TbDYRK. Finally, the direct substrate analysis was carried using procyclic form lysates in order to provide sufficient material for analysis, and so stage-specific differences in substrate proteins could contribute. Nonetheless, the comparison of both datasets revealed the presence of 45 common proteins exhibiting a change in phosphorylation regardless of the direction of change or the statistical value, with 2 proteins that had a statistical change in phosphorylation on the same peptides. The identified molecules showed an enrichment of RNA binding proteins, which is the predicted regulatory level of control for trypanosome development, given the emphasis in kinetoplastid parasites of post transcriptional regulation. Interestingly, conventional DYRK3 members in other eukaryotes have been identified as regulators of stress granule integrity, with the activity of the kinase influencing the dissolution of these membraneless organelles in order to control the response of cells to stress through regulation of the action of mTOR[45, 46]. Trypanosome differentiation similarly requires the regulation of TOR activity and the control of development through the dynamic association of predicted RNA binding proteins with stress granules, and TbDYRK - with features of DYRK2 and DYRK3 families - may functionally connect these regulatory components of trypanosome cell cycle exit and stumpy formation.

Genetic validation of some of the identified proteins in the stumpy differentiation pathway, by knock-out or conditional knock-down, revealed that TbDYRK acts on both the inhibitory and stimulatory arms of the differentiation control pathway. Indeed, knock-down of the hypothetical protein Tb927.4.2750 rendered the cells more sensitive to 8-pCPT-cAMP and knock-down of NOT5 led to an increase in cell differentiation at a lower parasitaemia. These observations suggest that the phosphorylation of these proteins by TbDYRK would inactivate their function, allowing the cells to differentiate. This identifies these proteins as so called ‘slender retainers’ that are required for the cells to remain as proliferative bloodstream forms. Conversely, the knock-out of the zinc finger protein ZC3H20 rendered cells unable to differentiate from slender to stumpy forms, indicating that its phosphorylation by TbDYRK would activate the protein to promote differentiation, a so called ‘stumpy inducer’. Consistent with this, the deletion of ZC3H20 rendered the cells unable to arrest in G1/G0 and this phenotype was reproduced when only the phosphosite targeted by TbDYRK was mutated, establishing a direct relationship between phosphorylation capacity and developmental competence. In combination this positions TbDYRK as a pivotal regulator of the trypanosome QS sensing pathway, directly acting on both differentiation activators and inhibitors through phosphorylation mediated control. Moreover, its effects on TbZC3H20 link the signal transduction cascade to post transcriptional regulation in the parasite, mRNAs regulated by this molecule being important in the parasite’s developmental events [34, 35].

## Materials and Methods

### Ethics statement

Animal experiments in this work were carried out in accordance with the local ethical approval requirements of the University of Edinburgh and the UK Home Office Animal (Scientific Procedures) Act (1986) under licence number 60/4373.

### Trypanosome culture, constructs and transfection

Pleomorphic *T. brucei brucei* EATRO 1125 AnTat1.1 90:13 cells [47] were used for the phenotypic analysis of the ectopic expression of mutants of the DYRK kinase and knock-down of the potential substrates in bloodstream forms. The *T. brucei brucei* EATRO 1125 AnTat1.1 90:13 cell line expresses the T7 RNA polymerase and tetracycline repressor protein. Pleomorphic *T. brucei brucei* EATRO 1125 AnTat1.1 cells were transfected by the plasmid J1339 (kindly provided by Dr. Jack Sunter), that contains the T7 RNA polymerase and the CRISPR/Cas9.

Slender forms were either grown *in-vitro* or harvested from MF1 female mice at 3 days post infection. Stumpy forms were harvested from MF1 female mice between 5- or 7- days post infection. All bloodstream form cell lines were grown *in-vitro* in HMI-9 at 37°C in 5% CO2. In vitro differentiation into ‘stumpy-like’ forms was performed for 96h using 100 µM of the cell permeable cyclic AMP analogue, 8-pCPT-cAMP, purchased from Sigma-Aldrich (United Kingdom) [9].

Ectopic expression constructs were generated using the pDex577-Y vector, that integrates in to the 177bp repeat mini-chromosome region. The different bloodstream RNAi cell lines were generated using the stem loop vector pALC14 [48]. Both ectopic expression and knock down were initiated by the addition of 2 μg/mL of doxycycline in the culture medium.

Null mutant constructs of the gene TbDYRK were created by replacing the YFP and TY tags of the pEnT6B-Y and pEnT6P-Y vectors[49] with fragments of the 3’ and 5’UTRs of the target gene then integrated in the genome, as describe in [8].

CRISPR/Cas9 knock-out construct of Tb927.7.2660 was generated as described in [50] using the pPOTv7 plasmid and transfected in the *T. brucei* EATRO 1125 AnTat1.1 J1339 strain. Correct integration of the construct and the deletion of the targeted gene were verified with the pairs of primers MC016/018 + MC017/018 and MC016/075 respectively.

Endogenous mutation of the gene coding for the protein ZC3H20 was generated, in the J1339 strain, by cloning the mutated version (OL068/069) of the gene into the pPOTv6-BSD plasmid using primers OL066/067 and the restriction enzymes HindIII/ScaI, replacing the tagging cassette. Integration was then performed as described in [50]. The second allele was deleted using the pPOTv7-HYG plasmid as described in [50] (Primer list in Supplementary Table 1). Correct mutagenesis and deletion were assessed by sequencing (Primer list in Supplementary Table 1).

Pleomorph transfections were performed as described by [51]. Selection was applied by using the appropriate drugs: Geneticin (G418, 90:13 = 2.5 μg/ ml), Hygromycin (HYG, All strains = 0.5 μg/ ml), Puromycin (PURO, 90:13 = 0.25 μg/ ml, J1339 = 0.05 μg/ ml), Blasticidin (BSD, 90:13 = 10 μg/ ml, J1339 = 2 μg/ ml) and Phleomycin (BLE, 90:13 = 2.5 μg/ ml).

#### pDEX577-Y

pDEX577_Tb927.10.15020::YFP-TY was generated by Gibson cloning as described in [8]. Site direct mutagenesis (Primer list in Supplementary Table 1) was then performed to generate the different mutants and their fidelity confirmed by sequencing (Primer list in supplementary table 1). Plasmids were linearized with NotI, prior to transfection into the *T. brucei* EATRO 1125 AnTat1.1 90:13 strain.

#### pALC14

RNAi target gene fragments were selected based on the default settings of the RNAit software [52]. Fragments were amplified using the pairs of primers indicated in the primer list in Supplementary Table 1 and cloned into pCR™-Blunt II-TOPO^®^(Invitrogen), prior to sequencing. Resulting constructs were then digested by HindIII/XbaI and the extracted fragments cloned into pALC14 plasmid opened by the same enzymes. The second round of cloning was performed by digestion by BamHI/XhoI, allowing the head-to-head arrangement of the 2 identical fragments into the pALC14, generating the stem loop [48]. Final plasmids were linearized with NotI, prior to transfection into the *T. brucei* AnTat1.1 90:13 strain.

#### pFASTBac1_TY-YFP-TY::Tb927.10.15020

Amplification of TY-YFP from pDEX577-Y was performed using the primer pair MC003/004 and cloned into pGEMTeasy (Promega), prior to sequencing (MC007/MC015). Next, the PstI restriction site into the YFP gene was removed by site directed mutagenesis using MC043/044 and the construct was linearised by the enzymes BssHII/PstI to allow the ligation of the hybridised primers MC005/006, for the integration of the second TY tag. The final tag TY-YFP-TY was then extracted from pGEMTeasy by BamHI/PstI and cloned into the pFASTBac1 vector (Gibco) previously digested by the same set of enzymes, to generate the pFASTBac1_TY-YFP-TY plasmid, allowing tagging in N-terminal or C-terminal. Cloning at the N-terminal end was performed by amplification of the gene TbDYRK with the primers MC001/002, subcloned into pGEMTeasy for sequencing (MC007/010/011/012/013/014/015), and digestion by BssHII/PstI to generate the final pFASTBac1_TY-YFP-TY::Tb927.10.15020. This final plasmid was then transformed into Bac10 bacteria (Gibco) to generate the baculovirus, necessary for the infection of insect cells, according to manufacturer instructions.

Site direct mutagenesis were then performed to generate the different mutants and then verified by sequencing.

### Protein expression and purification

Expression and purification of the recombinant tagged TY-YFP-TY:: TbDYRK was performed using the SF9 insect cell (Gibco) in SF900 II serum free medium, according to manufacturer’s instructions. Lysis of infected SF9, by the baculovirus allowing expression of the kinase, was performed with RIPA buffer (25mM Tris pH7-8, 150mM NaCl, 0.1%SDS, 0.5% Sodium deoxycholate, 1% Triton X-100, Protease inhibitor cocktail Roche mini EDTA-free, Benzonase at 25U/mL) and incubated 30 min on ice. After sonication, samples were clarified and the supernatant incubated with the αBB2 antibody [53]. Immunoprecipitation was then performed using the magnetic Dyneabeads protein G (Invitrogen), according to manufacturer’s instructions. Elution was carried out with 25mM Tris pH7.4.

#### Immunofluorescence

Cell cycle analysis and PAD1 protein expression analysis were carried out by staining ice-cold methanol fixed cells with 4’,6-diamidino-2-phenylindole (DAPI) (100 ng/ ml) and an anti-PAD1 antibody as previously described [22].

#### Protein visualisation and western blotting

Protein samples were boiled for 5 min in Laemmli loading buffer (except PAD1 samples, that remained unboiled), separated by SDS–PAGE (NuPAGE gel 4–12% Bis-Tris, Invitrogen) and visualized either by Coomassie staining or SYPRO Ruby Protein Gel Stain (Invitrogen) using a Typhoon 9400 scanner (Amersham Biosciences) with lex = 457 nm and lem = 610 nm. Alternatively, proteins were separated by SDS–PAGE on NuPAGE 4–12% Bis-Tris gels and blotted onto polyvinylidene difluoride (PVDF) membranes (Pierce). After blocking with Odyssey Blocking buffer for at least 30 min at room temperature (RT), membranes were incubated with antibodies 1- to 3-h at RT or overnight at 4°C under agitation in 2% BSA in TBS-T (0.1% Tween in TBS). The primary antibodies were used at the following dilutions: αPAD1 (1:1000); αEF1α (1:7000, Elongation Factor 1-alpha, Merck-Millipore); αBB2 (1:5,[53]); αGST (1:1000, Invitrogen); αThioP (1:1000, Thio-phosphate ester [51–8], Abcam). After 3 washes in TBS-T, proteins were visualised by incubating the membrane for 1h at RT with a secondary antibody conjugated to a fluorescent dye diluted 1:5000 in 50%Odyssey Blocking buffer/50% TBS-T. Finally, membranes were scanned using a LI-COR Odyssey imager system.

#### Northern blotting and transcriptome analysis

For northern blotting, RNA preparation and analysis were carried out as described by [8] with a hybridisation temperature of 62°C.

#### Kinase assay

The identification of a generic in-vitro substrate was performed using a ‘cold’ kinase assay with 10 μg dephosphorylated MBP, 40 μg histone H1, 1 μg histone cores H2A, H2B, H3, H4, 20 μg dephosphorylated casein, 1 μg β-Casein, or 1 μg of the recombinant *Mus musculus* Caspase 9 (Casp9, first 200 amino-acids - Cloud-Clone Corp, #RPA627Mu01) as substrates in kinase buffer at pH 7.5 (50 mM of MOPS pH 7.5, 100 mM NaCl, 10 mM MgCl2, 10 mM MnCl2) in 20 μl final volume and in the presence of 250 μM of adenosine-triphosphate (ATP)-γ -S [54]. After 30 min incubation at 37°C, the phospho-transferase reaction was stopped by an incubation for 10 min at 95°C, immediately followed by 2h of incubation in presence 5 mM PNBM at 20°C to initiate the alkylation reaction as described in [54]. The reaction was then stopped by adding Laemmli loading buffer. Reaction mixtures were separated by SDS–PAGE and transferred to PVDF membrane. Protein loading was revealed either by ponceau staining (substrate) or western blotting (kinase) using the αBB2 antibody. The phosphotransferase activity was revealed by western blotting using the αThioP antibody, as previously described [54].

The structure/function analysis was performed in a ‘hot’ kinase assay, using 5 percent of the TY-YFP-DYRK purified protein, incubated on a shaker for 25 min at 37°C with 0.5 μg of the Casp9 as substrate, 200 mM of ATP, 50 mM of MOPS pH 7.5, 100 mM NaCl, 10 mM MgCl2 and 1 mCi [γ-32P]-ATP (3000 Ci/mmol) in a final volume of 20 μl. The phosphotransferase reaction was then stopped by adding Laemmli loading buffer and boiling at 95°C for 5 minutes. Reaction mixtures were separated by SDS–PAGE and transferred to PVDF membrane. Protein loading was revealed either by ponceau staining (substrate) or western blotting (kinase) using the αBB2 antibody. ^32^P incorporation was monitored by exposing the membrane on an X-ray sensitive film (Roche) at −80°C. After exposure, the bands corresponding to Casp9 were excised from the PVDF membrane and Cherenkov radiation was quantified by a scintillation counter using the ^32^P program.

Kinase assays to identify substrates of the DYRK kinase was performed as follows: 15 percent of the TY-YFP-DYRK purified protein, incubated on a shaker for 30 min at 37°C with 1 mg of parasite lysate as substrate (see the phosphoproteomic section for the details), 200 mM of ATP, 50 mM of MOPS pH 7.5, 100 mM NaCl, 10 mM MgCl2. The kinase reaction was stopped by boiling samples for 5 min at 95°C.

#### RT-qPCR

RNA preparation was performed using the Qiagen RNA extraction kit, according to manufacturer’s instructions. 1 μg of RNA was treated with RQ1 RNase-free DNase (Promega) for 2 h at 37 °C before heat-inactivation. cDNA synthesis was performed using the SuperScript™ III Reverse Transcriptase (Invitrogen), according to manufacturer’s instructions, in presence of oligo(dT)_20_ (Invitrogen) and 500 ng of RNA. Real time PCR was performed using a LigthCycler® 96 (Roche). Oligonucleotides MC057/058 and MC059/060 amplified ∼120-150 bp fragments of TbDYRK or the YFP tag, respectively. Oligonucleotides MC055/056 [55], recognizing a fragment of GPI8, were used as an endogenous control for normalisation. PCRs were set up in triplicate, with each reaction containing 10 μL of Luna Universal qPCR Master Mix (New England BioLabs), 300 nM of each oligonucleotides, 5 μL of cDNA (diluted 1/10) in a final volume of 20 μL. PCR conditions were as follows: 1cycle of 50 °C for 2 min, 1 cycle of 95 °C for 10 min, followed by 50 cycles of 95 °C for 15 s and 58 °C for 1 min. Final melting curve was obtained by gradient increase temperature from 65°C to 95°C.

#### Phosphoproteomic analysis

The proteomic analysis comparing the phosphoproteome of the 2 strains *T. brucei* EATRO 1125 AnTat 1.1 90:13 WT or DYRK KO was performed exactly as described in [8]. Briefly, samples were extracted from 2 replicates for each cell line. For each, 2.7×10^8^ cells were lysed (4% SDS; 25 mM Tris(2-carboxyethyl)phosphine (TCEP) (Thermo); 50 mM N-ethylmaleimide (Thermo); 150 mM NaCl;1x PhosSTOP phosphatase inhibitor (Roche); 10 mM Na_2_HPO_4_ pH6) and then chloroform:methanol precipitated. Tryptic digestion was performed at 37°C with 8 μg trypsin (Pierce). Labelling and mass spectrometry were carried out at the FingerPrints Proteomics Facility at the University of Dundee. The samples were purified by solid phase extraction and then quantified by bicinchoninic acid assay prior to labeling with isobaric tandem mass tags (6-plex TMT). The samples were pooled and divided into fractions using hydrophilic interaction liquid chromatography. Statistical analysis of the fold change in phosphorylation between DYRK KO and *T. brucei* AnTat replicates was performed using the R package limma[56, 57].

The substrate identification analysis was performed with 2×10^9^ of procyclic cells. Lysates were generated using 4% SDS buffer (4% SDS, 25 mM Tris, 50 mM N-Ethylmaleimide (NEM), 150 mM NaCl, 10 mM Na2HPO4, Protease inhibitor cocktail Roche mini EDTA-free, 25U/mL Benzonase, pH6), incubated 30 min on ice, sonicated and heat inactivated for 5 min at 95°C. Next, samples were treated with Lambda Phosphatase (∼1U/μg of proteins) for 1h at 30°C, followed by another heat inactivation step of 5 min at 95°C. The kinase assay was then performed in 3 replicates with the active kinase NM or the inactive one (H866A), purified from insect cells, as described in the kinase assay section. To remove any trace of SDS, samples were precipitated with ice cold acetone, with a ratio of acetone/sample of 90%/10% and incubated at −20 overnight. The next day, samples were pelleted by centrifugation for 10 mins at 4°C, the supernatant discarded, and washed again with 1ml of ice-cold acetone and re-pelleted at 4 degrees. The following steps were performed at the proteomic facility of the University of Edinburgh. Dried pellets were resuspended in 8M urea and a protein assay performed (Bradford Biorad). One milligram of protein extract was digested (enough for two phosphopeptide enrichments). Protein denaturation and reduction was performed in 2M urea, 25mM ammonium bicarbonate and 5mM dithiothreitol (DTT).

Samples were kept at room temperature for 30 minutes before cysteine alkylation in 12.5mM iodoacetamide for 1h. Ten micrograms of trypsin were added and digestions were performed overnight at room temperature. Peptide extracts were then cleaned on an SPE reverse phase Bond Elut LMS cartridge, 25mg (Agilent). The samples were split into five hundred microgram aliquots and dried under low pressure (Thermo Jouan Speedvac) and stored at −20°C.

Samples were then resuspended with 25µl of 0.5M lactic acid/50% ACN and sonicated, prior phosphopeptide enrichment by adding 8µl of resin (100µg/µl of 10µm TiO beads in isopropanol - dried down). Following this, samples are mixed and incubated overnight at room temperature, with shaking. The sample/TiO2 mix was then transferred to a small filter and spun for 5min. 25µl of 0.5M lactic acid/50% ACN was added to the spin tip and then spun for 1min at 5000rpm in a microfuge. The resin washed with first 25µl 0.5M lactic acid/50% CAN and then with 200µl 80% ACN/0.1% TFA and spun for 1min at 5000rpm in a microfuge. Three additional washes were performed with 200µl 50% ACN/0.1% TFA and spin for 1min at 5000rpm, followed by 2 washes with 200µl 80% ACN/0.1% TFA and spin for 1min at 5000rpm. The first elution was performed as follows – 2 times with 50µl 50mM KH2PO4; the 2nd elution was as follows −50µl 2M Ammonia; and the 3rd elution comprised −50µl 80% ACN/0.1% TFA. The samples were dried under low pressure and reconstituted in 100 µl Buffer A (2%Acetonitrile in water 0.1%formic acid) before clean up. Clean-up of the eluted peptides was carried out with C18 membrane tips and then dried under low pressure. Once dried, the samples were reconstituted in 7µl 2% Formic Acid and filtered using a 0.45µm filter in preparation for MS.

Nano-ESI-HPLC-MS/MS analysis was performed using an on-line system consisting of a nano-pump (Dionex Ultimate 3000, Thermo-Fisher, UK) coupled to a QExactive instrument (Thermo-Fisher, UK) with a pre-column of 300 µm x 5 mm (Acclaim Pepmap, 5µm particle size) connected to a column of 75 µm x 50 cm (Acclaim Pepmap, 3 µm particle size). Samples were analysed on a 90 min gradient in data dependent analysis (1 survey scan at 70k resolution followed by the top 10 MS/MS). The gradient between solvent A (2%Acetonitrile in water 0.1%formic acid) and solvent B (80% acetonitrile-20% water and 0.1% formic acid) was as follows: 7min with buffer A, over 1 min increase to 4% buffer B, 57min increase to 25% buffer B, over 4min increase to 35%, over 1 min increase to 98% buffer B and continuation under those conditions for 9min, switch to 2% buffer B over 1 min and the column was conditioned for 10min under these final conditions. MS/MS Fragmentation was performed under Nitrogen gas using high energy collision dissociation in the HCD cell. Data was acquired using Xcalibur ver 3.1.66.10.

Data from MS/MS spectra was searched using MASCOT Versions 2.4 (Matrix Science Ltd, UK) against a *Trypanosoma brucei* database with maximum missed-cut value set to 2. The following features were used in all searches: i) variable methionine oxidation, and S/T/Y phosphorylation, ii) fixed cysteine carbamidomethylation, iii) precursor mass tolerance of 10 ppm, iv) MS/MS tolerance of 0.05 Da, v) significance threshold (p) below 0.05 (MudPIT scoring) and vi) final peptide score of 20. Progenesis (version 4 Nonlinear Dynamics, UK) was used for LC-MS label-free quantitation. Only MS/MS peaks with a charge of 2+, 3+ or 4+ were considered for the total number of ‘Feature’ (signal at one particular retention time and m/z) and only the five most intense spectra per ‘Feature’ were included.

### Bioinformatics approaches

DYRK orthologues of *Leishmania spp* and *Trypanosoma spp* genes and protein sequences were retrieved from the web database TriTrypDB (http://tritrypdb.org/tritrypdb/)[58]. Homology searches were carried out using BLAST with the default BLOSUM-62 substitution matrix [59], and pattern recognition analysis using the program PRATT v2.1 [60]. Multiple sequence alignments were performed using the built-in algorithm ClustalXv2. Additional sequence analyses were carried out using the Jalview program [61]. Statistical analysis and data plotting were performed using Rstudio software (http://www.rstudio.org/.rstudio.org/) and R language (R Development Core Team (2005). R: A language and environment for statistical computing. R Foundation for Statistical Computing, Vienna, Austria. ISBN 3-900051-07-0, URL: http://www.R-project.org).

### Statistical analyses

For the analysis of phenotypes, three to five animals per treatment were routinely used, with pilot and independent replicates confirming observed responses. With effects sizes similar to those previously observed for RNAi mediated loss of developmental competence (0.637 to 1.804; e.g.[9]) a sample size of three to five animals per group (+ or − DOX), or a total of six to ten, allows 80% power for test genes. Data were examined before analysis to ensure normality and that no transformations were required. Global proteomic data analyses were carried out using limma with a moderated t-test. Phosphoproteomic of the substrates of NM and the H866A mutant kinase in cell lysates was carried out using a standard t-test. *P* values of less than 0.05 were considered statistically significant. Blinding was not carried out.

## Acknowledgements

This work was funded by a Marie Sklodowska Curie postdoctoral fellowship to MC (proposal number 65470) and a Wellcome Investigator award (103740/Z14/Z) and Royal Society Research merit award (WM140045) to KRM.

## Supplementary Data

**Supplementary Figure S1:**
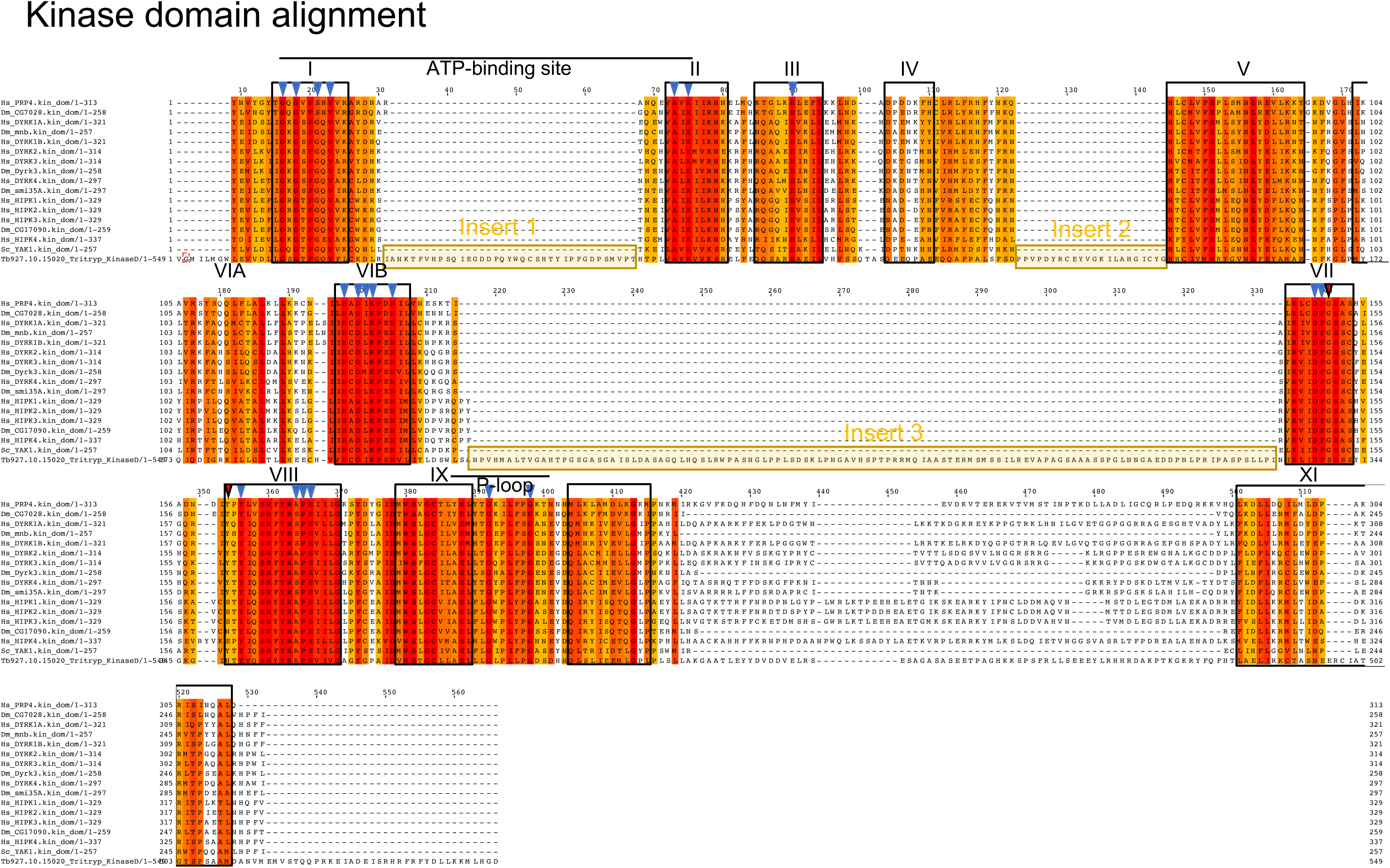
Multiple sequence alignment of the kinase core of TbDYRK (lower sequence) with other DYRKs from other species. Gradient colours indicate percentage of similarity, Latin numbering indicate the XII characteristic kinase domains, blue arrow heads indicate highly conserved residues, red arrow heads indicates usually highly conserved residues that are different in TbDYRK, yellow boxes highlight the 3 inserts.

**Supplementary Figure S2:**
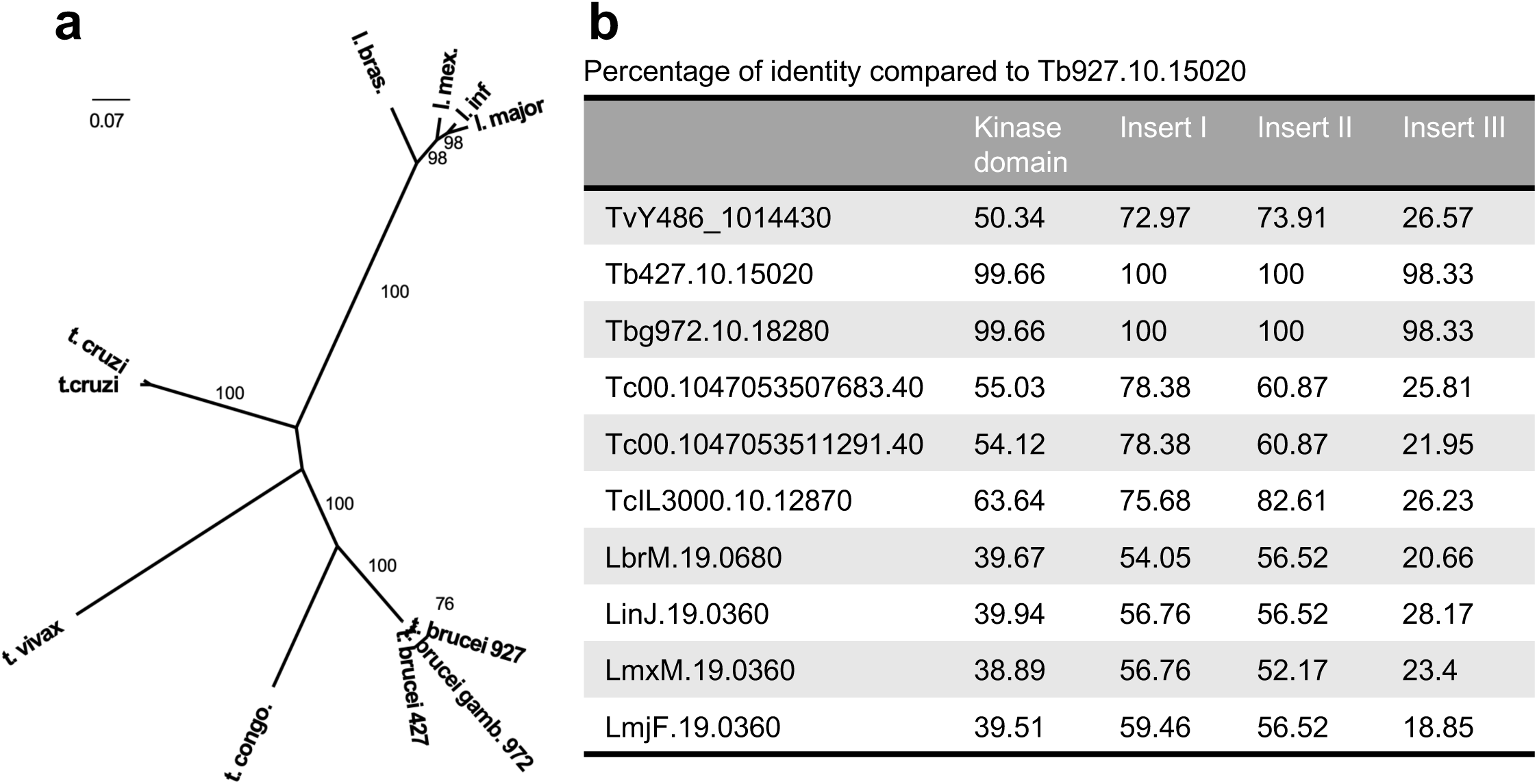
**a.** Molecular Phylogenetic analysis by the Maximum Likelihood method. The evolutionary history was inferred by using the Maximum Likelihood method based on the Whelan And Goldman + Freq. model [62]. The tree with the highest log likelihood (−6239.9546) is shown. The percentage of trees in which the associated taxa clustered together is shown next to the branches. Initial tree(s) for the heuristic search were obtained by applying the Neighbour-Joining method to a matrix of pairwise distances estimated using a JTT model. The tree is drawn to scale, with branch lengths measured in the number of substitutions per site. The analysis involved 11 amino acid sequences. All positions with less than 95% site coverage were eliminated. That is, fewer than 5% alignment gaps, missing data, and ambiguous bases were allowed at any position. There were a total of 542 positions in the final dataset. Evolutionary analyses were conducted in MEGA7 [63]. Bootstrap proportions higher than 70% are shown at internal nodes. **b.** Percentage of identity of different domains of the orthologues of the TbDYRK from other kinetoplastid species.

**Supplementary Figure S3:**
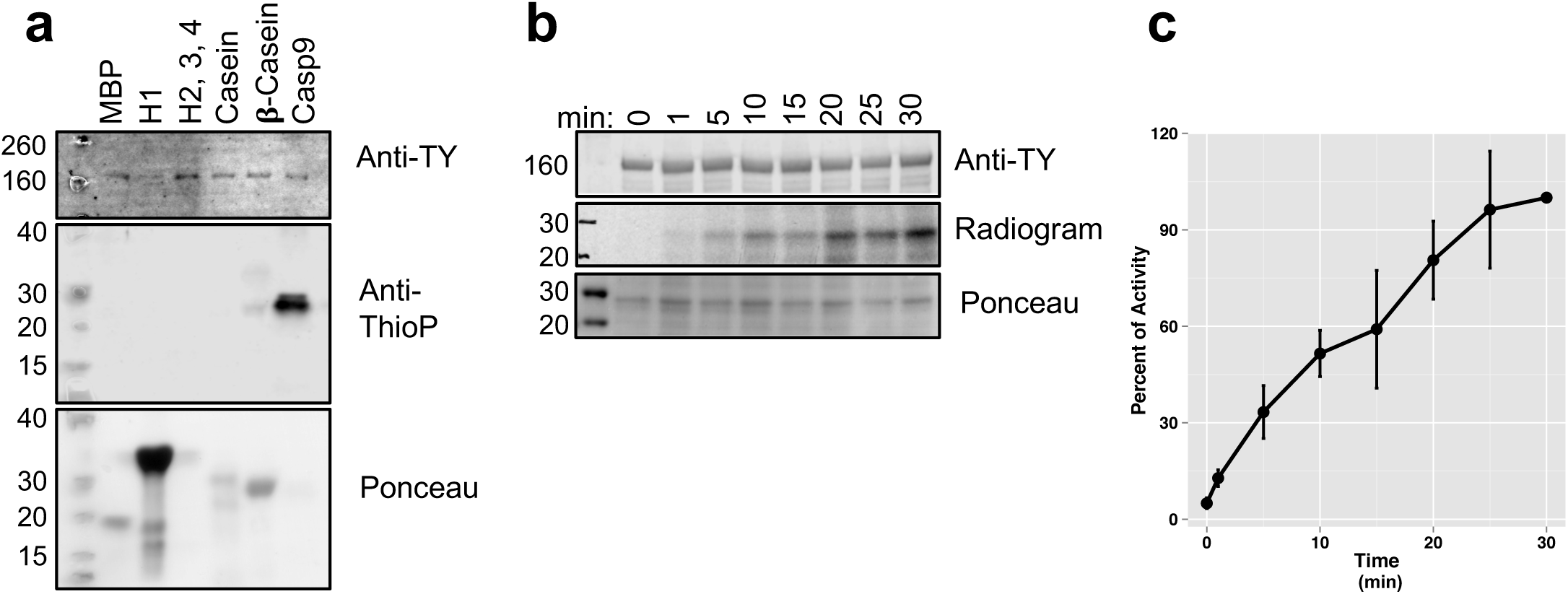
**a.** In vitro ‘cold’ kinase assay using the indicated proteins as potential generic substrates of the active kinase NM purified from insect cells. The kinase assay was revealed by Western blotting using the antibody anti-TY (revealing the loading of the kinase), the antibody anti-thiophosphate (anti-ThioP, revealing the phosphorylated substrates) and the ponceau staining to reveal the loading of the substrates. **b.** Kinetic analysis of the phosphorylation of the generic substrate Casp9 revealed by hot kinase assay. The radiogram presented in B is representative of 3 independent experiments. g. Quantification of the Cherenkov radiation signal of 3 independent experiments of the kinetic analysis presented in b. error bars=SEM

**Supplementary Figure S4:**
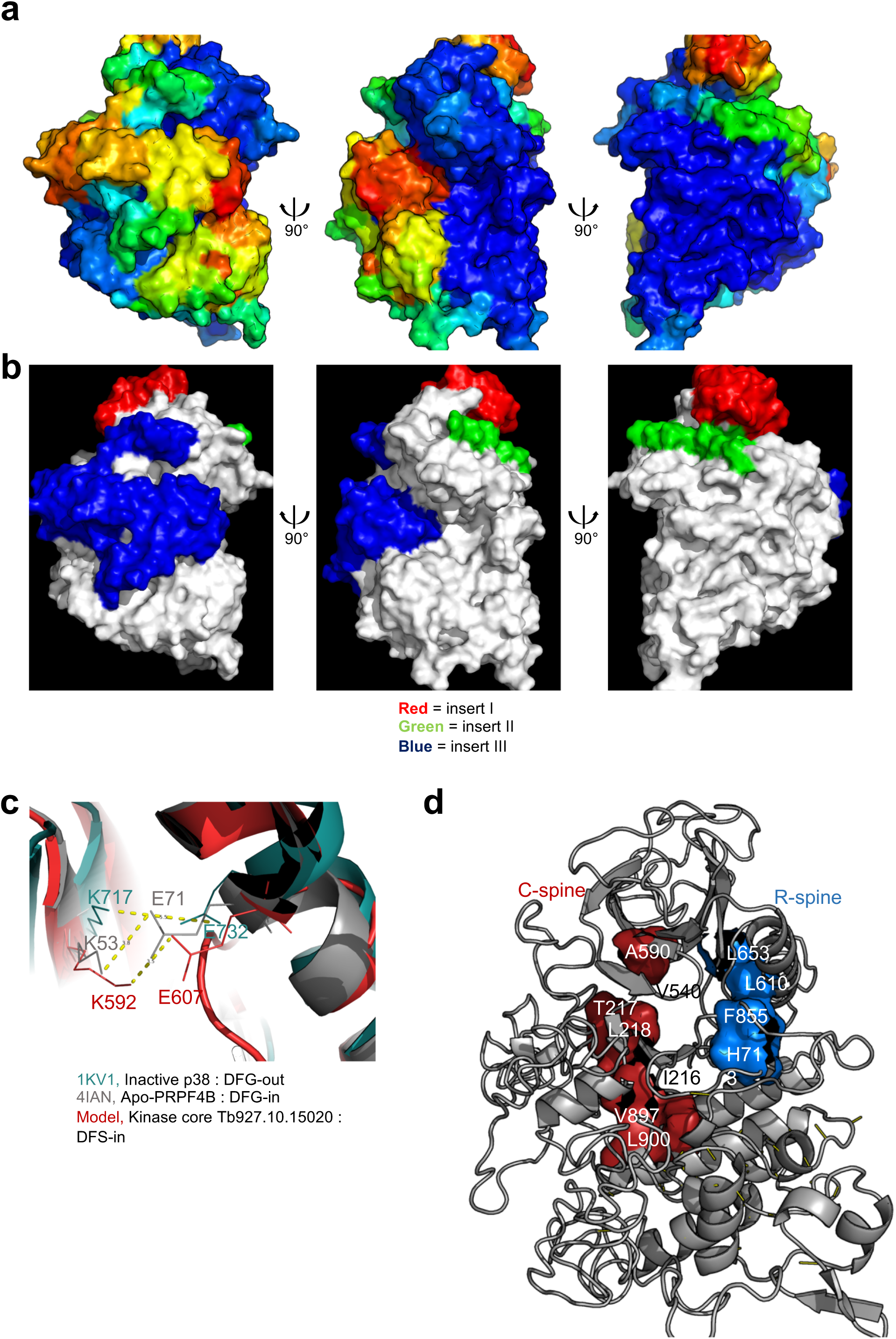
Analysis of cell viability by alamar blue of parasites expressing the ectopic DYRK NM / H866A / ΔII in response to cAMP treatment. Results have been obtained from 3 independent replicates and normalised by the -cAMP +Dox condition for each cell line, error bars=SD

**Supplementary Figure S5:**
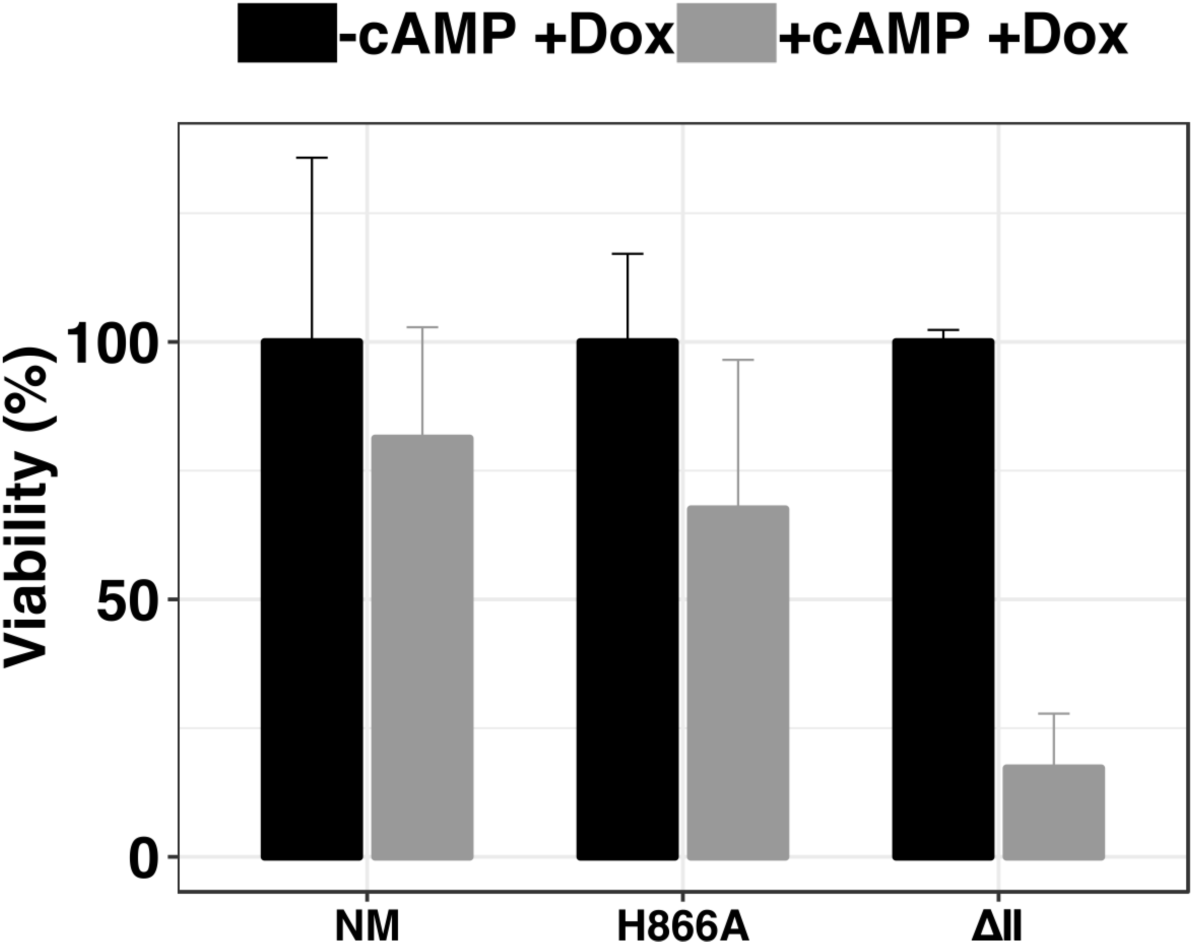
Modelling of the kinase core of TbDYRK. **a.** Visualisation of the b-factor score on the model obtained from the i-TASSER server. Blue = high confidence, red = low confidence. Three views rotated on the y axis by 90° are presented, with the left panel centred on the inserts, the middle panel on the ATP binding pocket and the right panel on the activation loop. **b.** Visualisation of the 3 inserts on the model obtained from the i-TASSER server. Red = insert I, green = insert II and blue = insert III. Three views are presented, centred and rotated as previously described in **a**. **c.** Superposition of the model to crystal structures in active conformation (PRPF4-B), and inactive conformation (p38), highlighting the distance between the lysine of the β3 strand of the N-lobe and the glutamic acid of the αC-helix for the close conformation of the ATP binding pocket. **d.** Cartoon representation of the kinase core of the model of TbDYRK highlighting the alignment of the non-sequential residues forming the C- and R-spines, as tipically observed in active conformations.

**Supplementary Figure S6:**
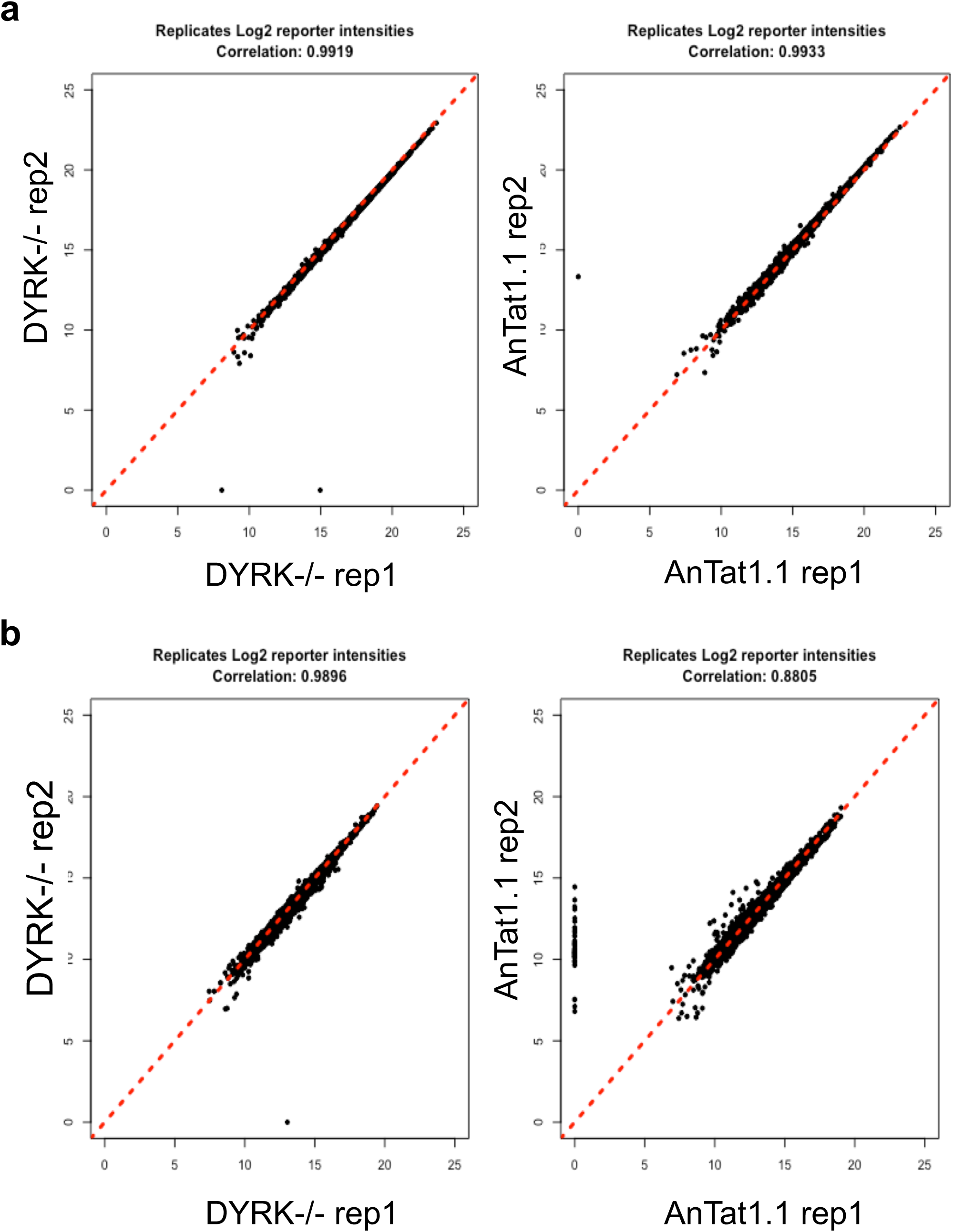
Correlation plots between replicates of the phosphoproteomic analysis comparing DYRK-/-cells to WT AnTat1.1 cells. **a**, Correlation plots of proteins, **b**, Correlation plots of phosphopeptides.

**Supplementary Figure S7:**
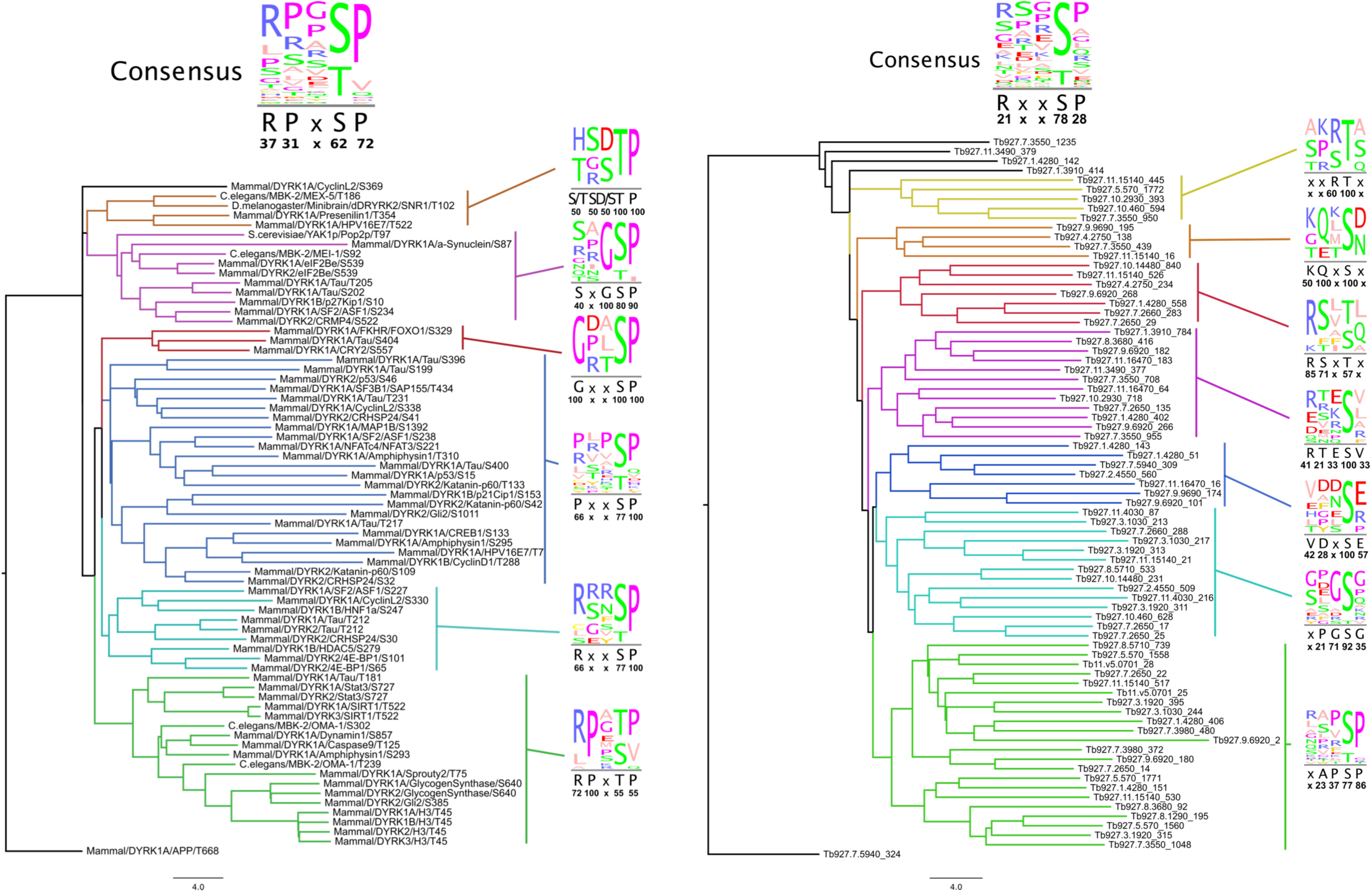
Substrate motif identification. Substrates from both Datasets: NMvsH866A (right panel) and WTvsKO (left panel) with |FC|>1.5. Trees generated by neighbour joining using BLOSUM62, consensus filtered for 20% appearance of the residue. Numbers below consensus motifs represent percentage of appearance of the corresponding residue.

**Supplementary Figure S8:**
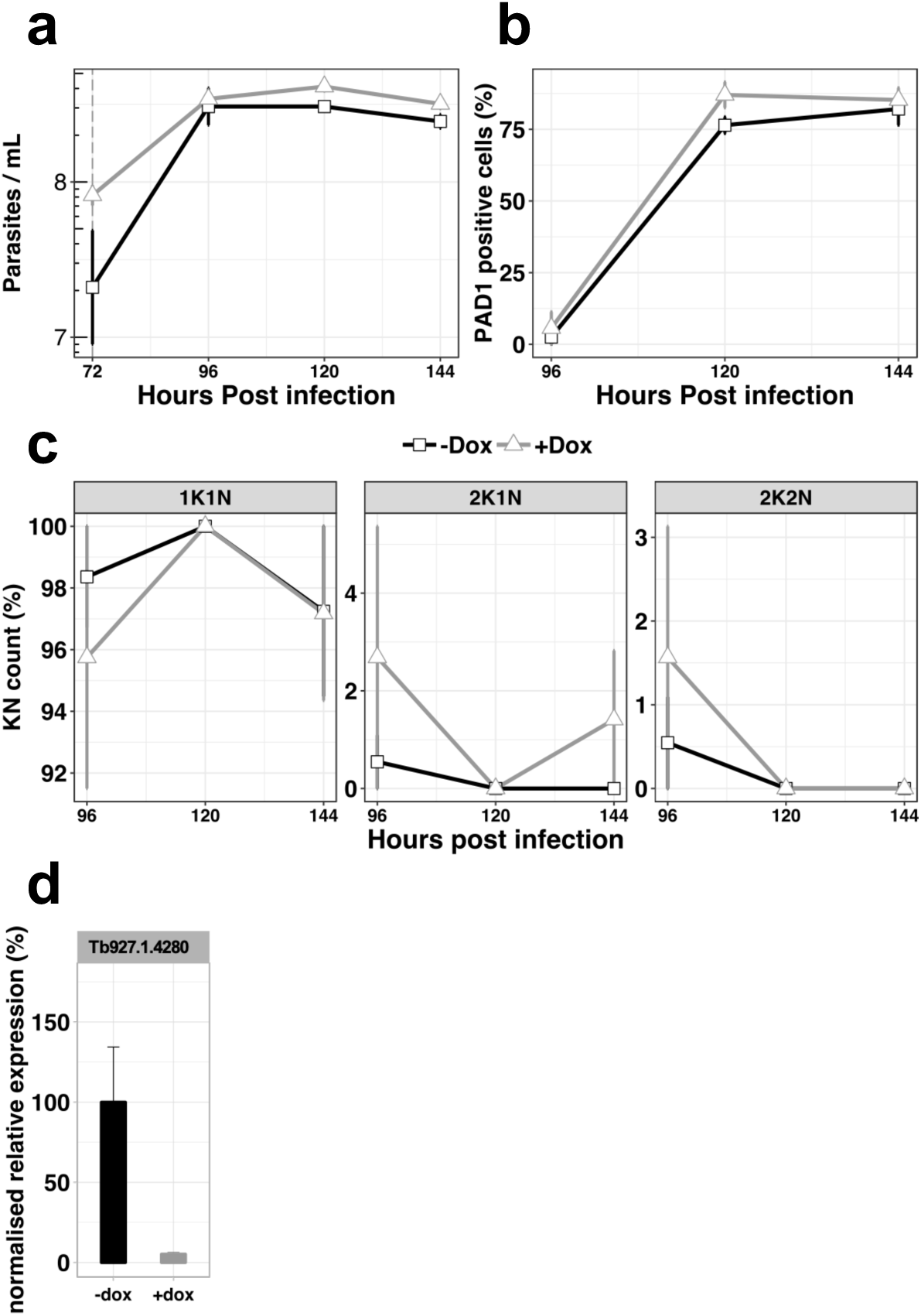
*In vivo* phenotypic analysis of the RNAi knocked-down cell line for Tb927.1.4280, uninduced (-dox, black line) or induced with doxycycline at 72h post infection (+dox, grey line). **a,** Parasitaemia, **b,** Percentage of PAD1 positives cells, **c**, Percentage of KN count, **d**, RT-qPCR determining the relative expression of the gene of interested. n=3, error bars=SD.

**Supplementary Figure S9:**
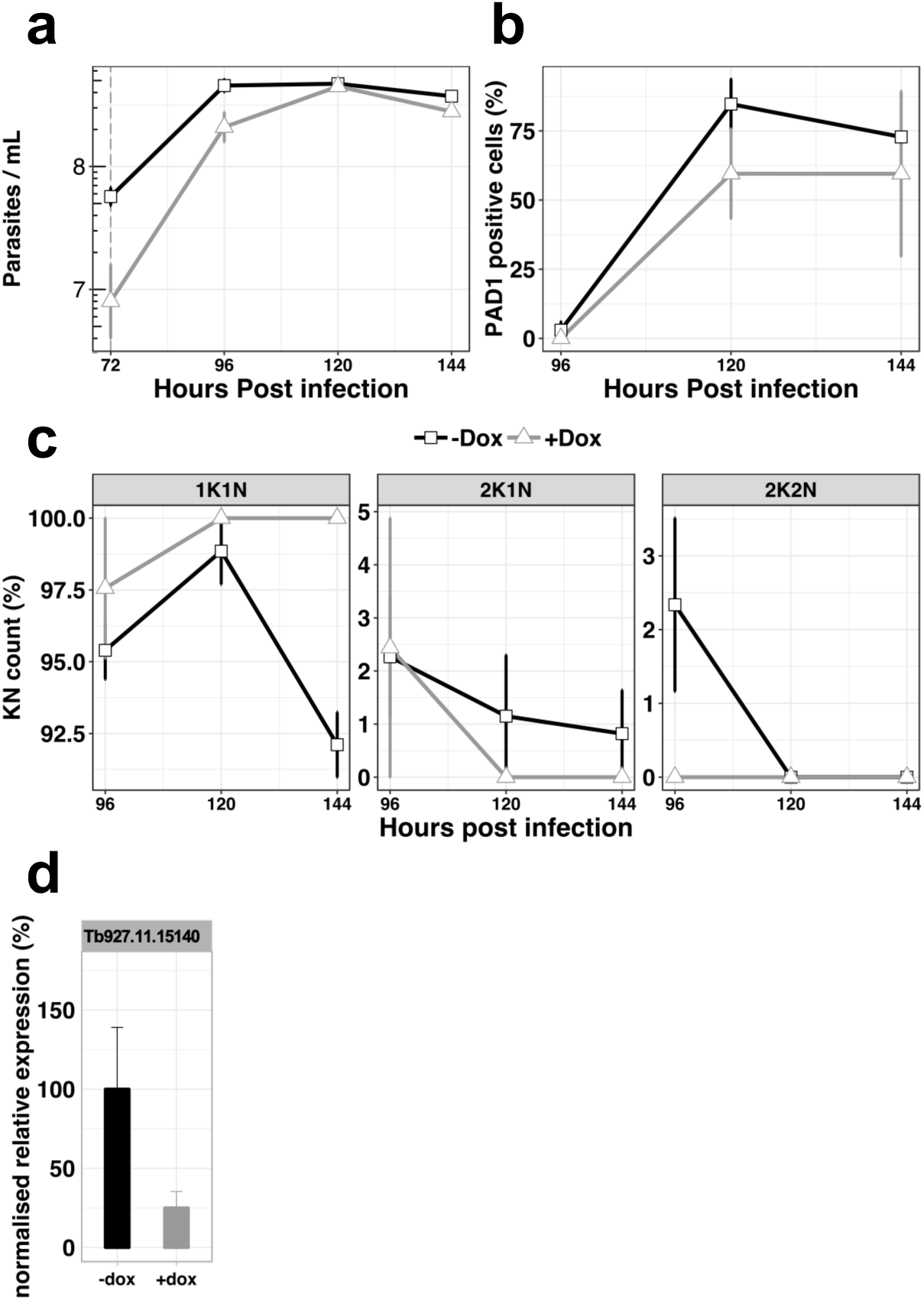
*In vivo* phenotypic analysis of the knocked-down cell line for Tb927.11.15140, uninduced (-dox, black line) or induced with doxycycline at 72h post infection (+dox, grey line). **a**, Parasitaemia; **b**, Percentage of PAD1 positives cells, **c**, Percentage of KN count, **d**, RT-qPCR determining the relative expression of the gene of interested. n=3 error bars=SD.

**Supplementary Figure S10:**
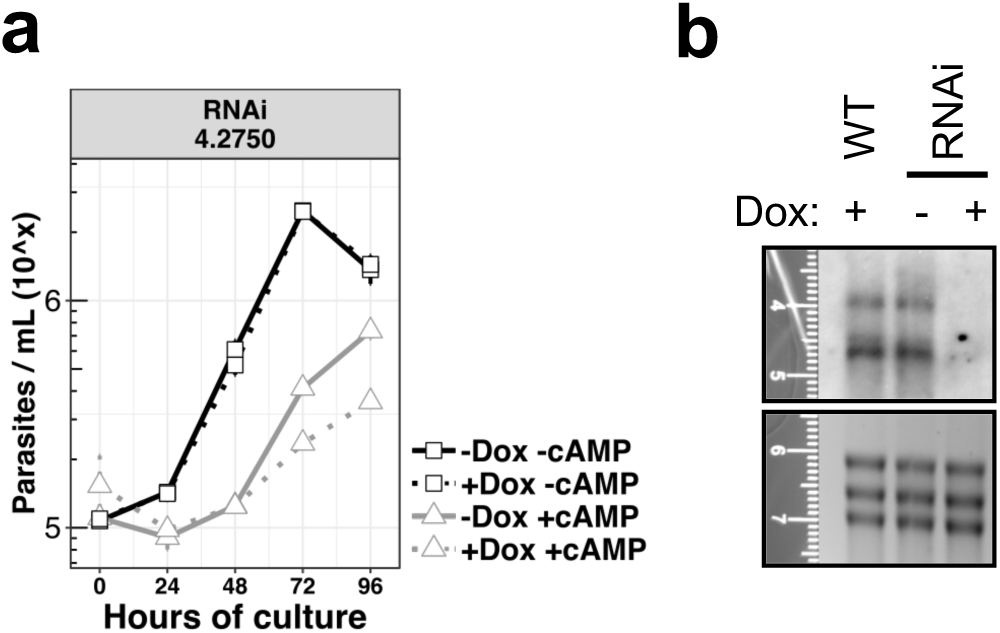
*In vitro* phenotypic analysis of the knocked-down cell line for gene Tb927.4.2750, uninduced (-dox) or induced with doxycycline (+dox), and treated or not with the cell permeable cAMP. **a**, Growth curves, n=3 error bars=SD; **b**, Northern blot analysis representative of the 3 replicates probing the gene Tb927.4.2750 in WT cells or in the knock-down cell line (RNAi) induced or not with doxycycline.

**Supplementary Figure S11:**
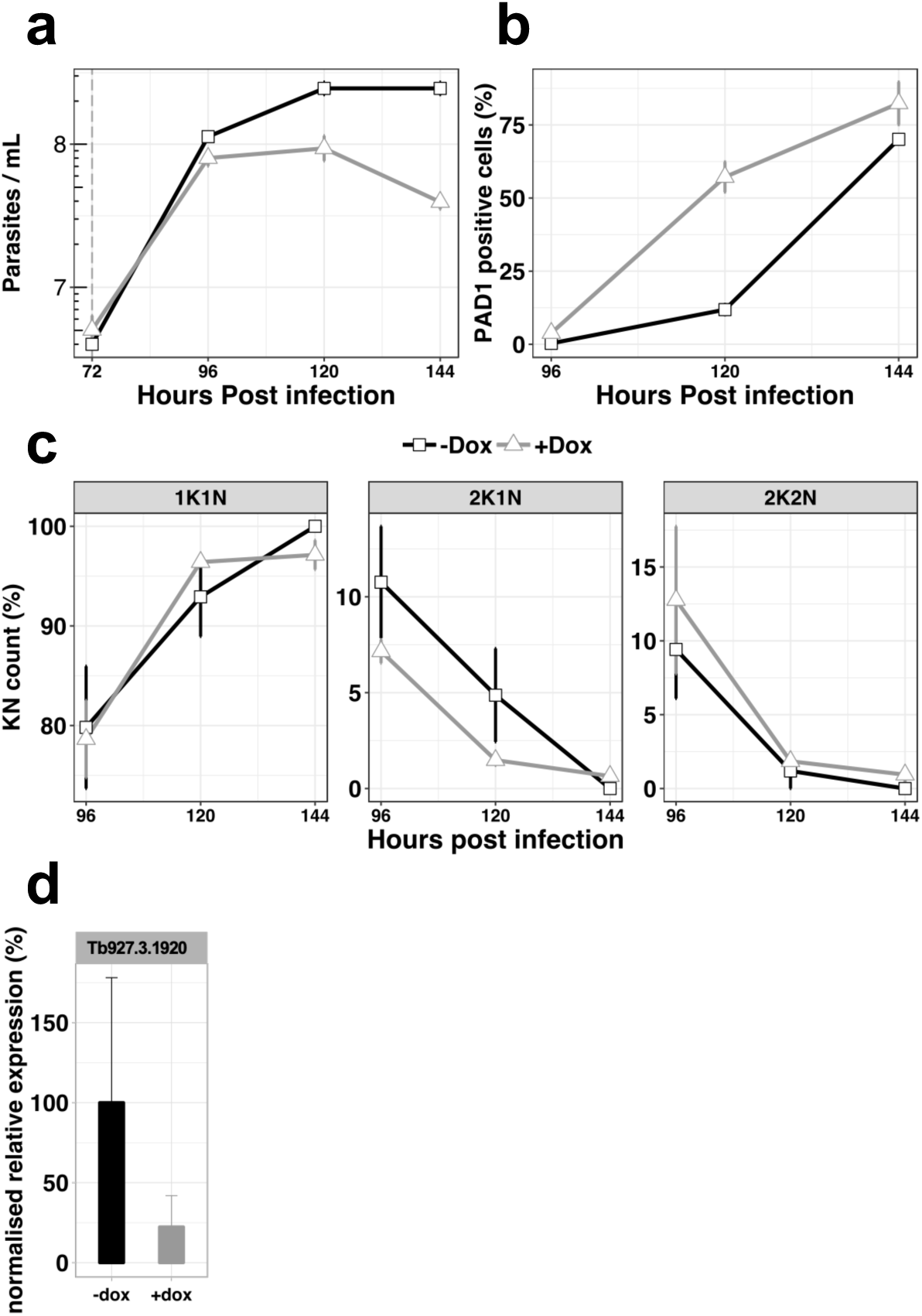
*In vivo* phenotypic analysis of the knocked-down cell line for gene Tb927.3.1920, uninduced (-dox, black line) or induced with doxycycline at 72h post infection (+dox, grey line). **a**, Parasitaemia, n=3, error bars=SD; **b**, Percentage of PAD1 positive cells, n=3, error bars=SD, c) Percentage of KN count, n=3 error bars=SD d) RT-qPCR determining the relative expression of the gene of interested. n=3, error bars=SD.

**Supplementary Figure S12.**
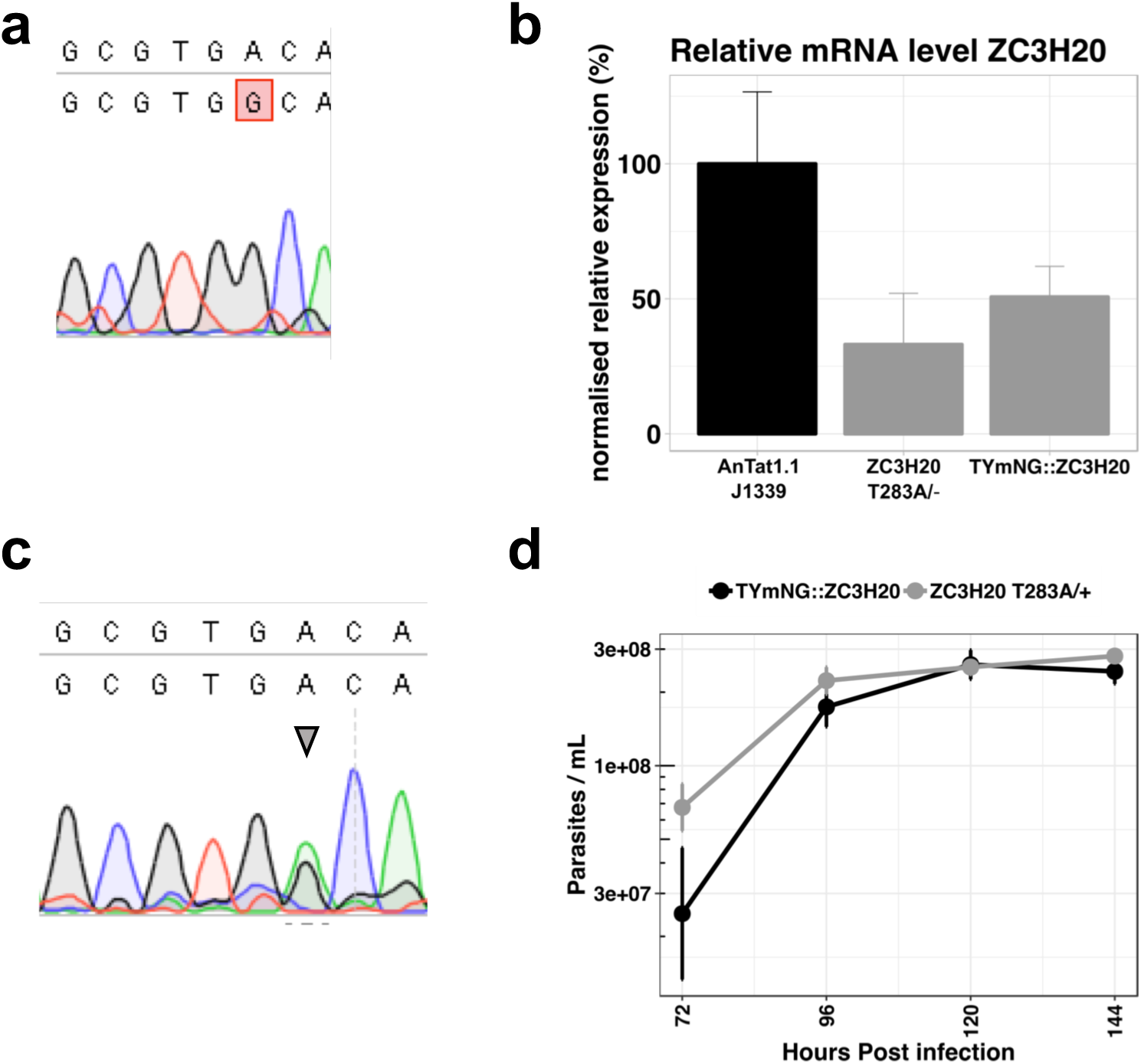
Endogenous mutation ZC3H20 T283A. **a.** Sequencing results of the gene Tb927.7.2660 coding for the protein ZC3H20 presenting the single allele mutation A to G generating the T283A. **b.** RT-qPCR analysis revealing the mRNA expression level of ZC3H20 in the single allele ZC3H20 ^T283A/-^ compared to the parental cell line and a line with a single allele of ZC3H20 replaced by mNeon Green (TYmNG::ZC3H20). n=3 error bars=SEM **c.** Sequencing results of the gene Tb927.7.2660 coding for the protein ZC3H20 presenting both the single allele mutation A to G generating the T283A and the endogenous WT allele (arrow head). **d.** *In vivo* parasitaemia of cell line ZC3H20 T283A/+ (grey line) compared to the parental cell line WT (black line). n=3 error bars=SD

**Supplementary Table 1:** List of primers used in the study.

**Supplementary Table 2:** Results of the phosphoproteomic analysis comparing DYRK-/- cells to WT AnTat1.1cells. Sheet 1 KOvsWT_|FC|>1.5_p<0.05 presents the peptides with an absolute value of FC>1.5 and a P-value<0.05. Sheet 2 GOenrichDownKO indicate the GO term enrichment of biological functions of proteins presenting a down regulation of phosphorylation in the DYRK-/- cell line. Sheet 3 GOenrichUpKO indicate the GO term enrichment of biological functions of proteins presenting an up regulation of phosphorylation in the DYRK-/- cell line. Yellow colour indicates the proteins for which further analysis have been performed.

**Supplementary Table 3:** Results of the phosphoproteomic analysis the phosphorylation of cell lysates by the active DYRK non-mutated NM to the inactive mutant H866A. Sheet 1 NMvsH866A_|FC|>1.5_p<0.05 presents the peptides with an absolute value of FC>1.5 and a P-value<0.05. Sheet 2 GOenrichUpNM indicate the GO term enrichment of biological functions of proteins presenting a phosphorylation in presence of the NM. Yellow colour indicates the proteins for which further analysis have been performed.

**Supplementary Table 4:** List of proteins (sheet 1) and peptides (sheet 2) common in both datasets with an absolute value of FC>1.5. Green colour indicates the statically significant protein/peptides (P-value<0.05).

